# Bursts in biosynthetic gene cluster transcription are accompanied by surges of natural compound production in the myxobacterium *Sorangium* sp

**DOI:** 10.1101/2022.11.24.517636

**Authors:** Judith Boldt, Laima Lukoševičiūtė, Chengzhang Fu, Matthias Steglich, Boyke Bunk, Vera Junker, Aileen Gollasch, Birte Trunkwalter, Kathrin I. Mohr, Michael Beckstette, Joachim Wink, Jörg Overmann, Rolf Müller, Ulrich Nübel

## Abstract

We have investigated the time course of genome-wide transcription in the myxobacterium *Sorangium* sp. So ce836 in relation to its production of natural compounds. Time-resolved RNA sequencing revealed that core biosynthesis genes from 48 biosynthetic gene clusters (BGCs; 92% of all BGCs encoded in the genome) were actively transcribed at specific time points in a batch culture. The majority (80%) of polyketide synthase and non-ribosomal peptide synthetase genes displayed distinct peaks of transcription during exponential bacterial growth. Strikingly, these bursts in BGC transcriptional activity were associated with surges in the production of known natural compounds, indicating that their biosynthesis was critially regulated at the transcriptional level. In contrast, BGC read counts from single time points had limited predictive value about biosynthetic activity, since transcription levels varied >100-fold among BGCs with detected natural products. Taken together, our time-course data provide unique insights into the dynamics of natural compound biosynthesis and its regulation in a wild-type myxobacterium, challenging the commonly cited notion of preferential BGC expression under nutrient-limited conditions. The close association observed between BGC transcription and compound production warrants additional efforts to develop genetic engineering tools for myxobacterial producer strains, to boost compound yields by manipulating transcriptional activity.

## Introduction

Natural products are important sources of biologically active molecules for the discovery of novel drugs (Newman & Cragg, 2020). In particular, the majority (71%) of small-molecule antibiotics approved for clinical use since the 1980s are derived from natural products (Newman & Cragg, 2020). Two-thirds of all naturally derived antibiotic compounds originate from actinobacteria (Barka et al., 2016), and consequently, their physiology and molecular biology have been studied intensely for several decades (van der Heul et al., 2018). In recent years, tremendous bacterial genome data deposited in public databases revealed that many bacteria from other phyla also have great potential for the synthesis of natural products (Gavriilidou et al., 2022). One of these groups of organisms are myxobacteria, which produce diverse natural products with unique structural features and bioactivities (Herrmann et al., 2017). Among the myxobacteria, the most prolific source of biomedically useful metabolites is as yet the genus *Sorangium* (Gerth et al., 2003; Hoffmann et al., 2018), which comprises at least eight distinct species (Mohr et al., 2018). Natural products from *Sorangium* spp. include for example, the antitumor compound epothilone (Altmann et al., 2009), the antibacterial and anti-biofilm compound carolacton (Jansen et al., 2010), and the antifungal compound icumazole (Barbier et al., 2012).

Even though natural compounds display enormous structural diversity, they originate from limited classes of biosynthetic pathways with common features, providing the basis for their classification into polyketides (PKs), non-ribosomal peptides (NRPs), ribosomally synthesized and post-translationally modified peptides (RiPPs), terpenoids and few others. The genes involved in a specific biosynthesis pathway are commonly arranged at adjacent positions on the bacterial producer’s chromosome, in so-called biosynthetic gene clusters (BGCs) (Fischbach & Walsh, 2006). The bioinformatic detection of BGCs in DNA sequence data with the software tool antiSMASH uses a rule-based approach and relies on the identification of core biosynthetic genes, for example, those encoding PK synthases or NRP synthetases, based on their conserved modular structures (Blin et al., 2021). Accessory genes surrounding the core biosynthetic genes within a predicted BGC region are annotated as additional biosynthetic, regulatory, or transport-related genes, respectively, relying on secondary metabolite clusters of orthologous groups of proteins (smCOGs) (Medema et al., 2011). Co-localized genes that do not fall into any of these categories are classified as ‘other genes’.

Systematic analyses of previously described natural products suggested the existence of large natural reservoirs of structural novelty still awaiting discovery (Gregory et al., 2019; Pye et al., 2017). This insight has been corroborated by genomic analyses of microbial producer strains, which have repeatedly documented the genetic potential for compound biosynthesis to vastly exceed the diversity of natural products detected in laboratory cultures (Aigle et al., 2014; Baltz, 2017; Jones et al., 2016). The majority of BGCs could not be matched to any detected product, and have thus frequently been interpreted as ‘silent’, i.e. not being expressed under standard laboratory conditions (Hoskisson & Seipke, 2020; Okada & Seyedsayamdost, 2017). Indeed, in actinobacteria the production of several PKs and NRPs has been demonstrated to require triggering by specific external cues (van der Heul et al., 2018). Notably, the depletion of phosphate (Martín, 2004; Thomas et al., 2012), nitrogen (Voelker & Altaba, 2001), or carbon sources (Sánchez et al., 2010), among other stress factors, was identified to stimulate the biosynthesis of antibiotics and other ‘secondary metabolites’ (van Wezel & McDowall, 2011; Wohlleben et al., 2017). In *Streptomyces* spp., the transcriptional activity of genes encoding enzymes for synthesizing natural compounds was mostly observed at later stages of batch cultivation, after nutrients had been depleted, whereas it was repressed during exponential bacterial growth (Nieselt et al., 2010; X.-M. Zhu et al., 2019). It is not clear, however, to what extent a generalization of this observation from actinobacteria is justified for other microorganisms, since, for example, myxobacteria produce several natural compounds preferably during growth in nutrient-rich media (Gerth et al., 1982; Kegler et al., 2006; Kunze et al., 1984; Rachid et al., 2007).

Investigations into the cellular regulation of BGC activities could facilitate control over the production of novel bioactive compounds, either by targeted molecular manipulation (Mungan et al., 2022; van der Heul et al., 2018) or by providing suitable extracellular cues (Okada & Seyedsayamdost, 2017). Transcriptional activities of BGCs and their regulation have been studied in *Streptomyces* (Hwang et al., 2019; Nieselt et al., 2010; X.-M. Zhu et al., 2019) and related actinobacteria (Tocchetti et al., 2015), and in a few other model organisms (Neubacher et al., 2020; Wu et al., 2020). Insights into genetic regulatory circuits of natural compound synthesis in myxobacteria are more limited at present. Most previous transcriptomic studies on myxobacteria were driven by an interest in the genetic regulation of their multicellular developmental program (McLoon et al., 2021; Müller et al., 2010; Muñoz-Dorado et al., 2019; Sharma et al., 2021) or changes during co-operative predation on other bacteria (Livingstone et al., 2018), focusing on the model species *Myxococcus xanthus*. In agreement with the triggering events for antibiotic production in actinobacteria, the transcription of genes related to the biosynthesis of some polyketides and terpenoids in agar plate cultures of *M. xanthus* was shown to be enhanced after UV irradiation, presumably as part of a cellular SOS response (Sheng et al., 2021). In contrast, an early study on *Sorangium* sp. So ce56 that applied quantitative reverse-transcription PCR, demonstrated that three core biosynthesis genes in the BGC for the macrocyclic compound chivosazol were maximally transcribed during exponential growth of the bacterium, coinciding with the production of chivosazoles (Kegler et al., 2006). In follow-up studies, Rachid *et al*. showed that transcription of the chivosazol BGC was under the control of the pleiotropic, positive regulator ChiR and the ammonia-responsive, negative regulator NtcA (Rachid et al., 2007, 2009). Furthermore, these authors succeeded in increasing chivosazol production by overexpressing *chiR* or inactivating the *ntcA* gene in the natural producer strain. Enhanced transcription of the BGC encoding epothilone biosynthesis in *Sorangium* sp. was achieved by co-cultivation with other *Sorangium* spp. strains (P. Li et al., 2013) and by targeted molecular manipulation of the BGC promoter (Ye et al., 2019). In both these reports, enhanced transcription of BGC genes was directly associated with increased epothilone production yields (P. Li et al., 2013; Ye et al., 2019). Activation of silent biosynthetic gene clusters in myxobacteria was achieved by manipulation of the endogenous promotor sequences resulting in the identification of completely novel natural products (Panter et al., 2018). These examples demonstrate that a more thorough understanding of BGC expression and its genetic regulation will improve our abilities to control natural compound biosynthesis, to induce novel BGCs, or increase compound production yields.

## Results

### Biosynthetic potential of the genome of *Sorangium* sp. So ce836

The So ce836 genome has a size of 14,555,308 bp and encodes 10,707 genes, including 10,573 protein-coding sequences. Analysis with the bioinformatic tool antiSMASH (Blin et al., 2021) predicted a total of 45 BGC regions. Our detailed analysis of the BGC regions showed that four of these likely constituted several BGCs (see below), resulting in a total of 52 BGCs. Among these BGCs, only six showed close similarity (i.e. above 60% amino acid similarity) to BGCs with known natural products reported in the MIBiG database (Suppl. Table S1) (Kautsar et al., 2019). Thirty BGCs encoded the syntheses of NRPs, PKs, or hybrids of these (Suppl. Table S1). Predicted BGCs encompassed a total of 414 genes, comprising 4% of all proteincoding sequences in So ce836, and 307 of these (74%; 81 core biosynthetic, 131 additional biosynthetic, 42 transport-related, 53 regulatory) were associated with the production of NRPs and PKs.

### Temporal transcription patterns in a batch culture

In a batch liquid culture incubated at 28°C with vigorous shaking, So ce836 grew with a minimum doubling time of 2.3 days (***Figure 1***). The exponential growth phase lasted from day 1 to day 8, followed by a stationary phase and a death phase (***Figure 1***).

**Figure 1.**
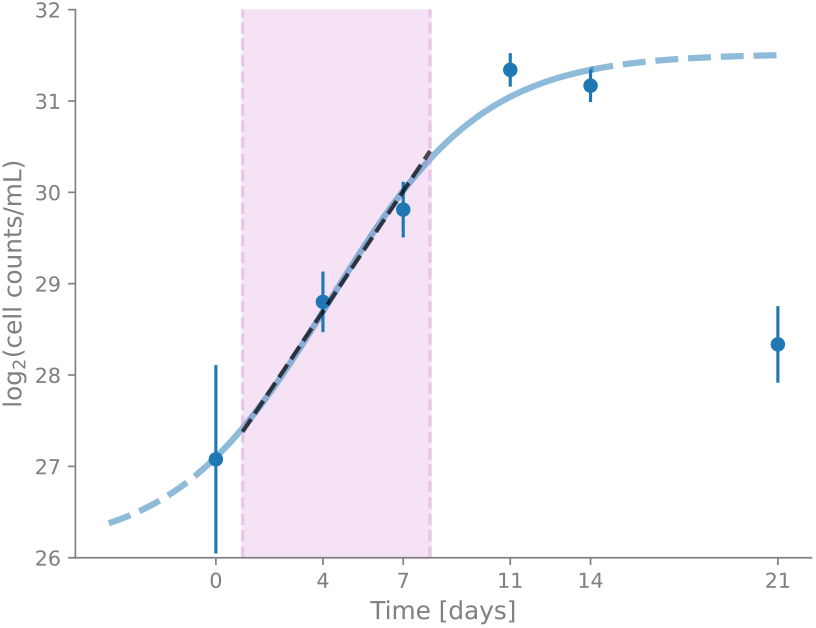
Growth curve of So ce836. The blue curve shows the growth of So ce836 over six time points (logistic curve *y* = *L*/(1 + *e*^-*k*\*(*x*–*x*0)^)+*b*, with *L* = 5.434, *k* = 0.347, *x*0 = 4.225, *b* = 26.083). The exponential phase (day 1 to day 8) is indicated by the pink area and black dashed line. Each data point represents mean cell counts per mL on a logarithmic scale (five biological replicates per growth time with ten technical replicates each). Error bars indicate the relative standard error of the mean.

We analyzed the time course of genome-wide transcription in the So ce836 cultures by sequencing RNA from samples collected at five different time points: day 4, day 7, day 11, day 14, and day 21. The transcription of 8,329 genes (78% of all genes) varied significantly over the time-course experiment, as indicated by a likelihood ratio test. These are referred to as ‘differentially expressed’ genes in the following text. In contrast, the transcription levels of 16% of all genes did not vary significantly between time points, and an additional 6% of genes were excluded from further analyses due to low read counts. To compare time courses of transcription between differentially expressed genes, we analysed deviations from the genes’ mean transcription levels at five time points. We found that the timing of transcription varied widely, and hierarchical clustering of these temporal patterns identified five transcription clusters (***Figure 2***). Genes in transcription cluster 2 (2,857 genes, 34% of differentially expressed genes) showed a tendency for maximal transcription on day 4, i.e. during early exponential growth (***Figure 2***). A majority of genes in transcription cluster 4 (1,423 genes, 17%) were maximally transcribed on day 7 of the experiment, i.e. also during exponential growth (***Figure 2***). In contrast, transcription cluster 5 (2,669 genes, 32%) contained genes that were strongly transcribed at the beginning of the stationary phase (day 11), and the transcription of genes in clusters 3 (395 genes, 5%) and 1 (985 genes, 12%) peaked on days 14 and 21, respectively (***Figure 2***). Hence, in summary, transcription clusters 2 and 4 mainly contained genes (totaling 40% of all genes) that reached their maximum transcription during exponential growth, whereas most genes in the other three transcription clusters (1, 3, and 5; 38% of all genes) were maximally transcribed in later growth phases (***Figure 2***).

**Figure 2.**
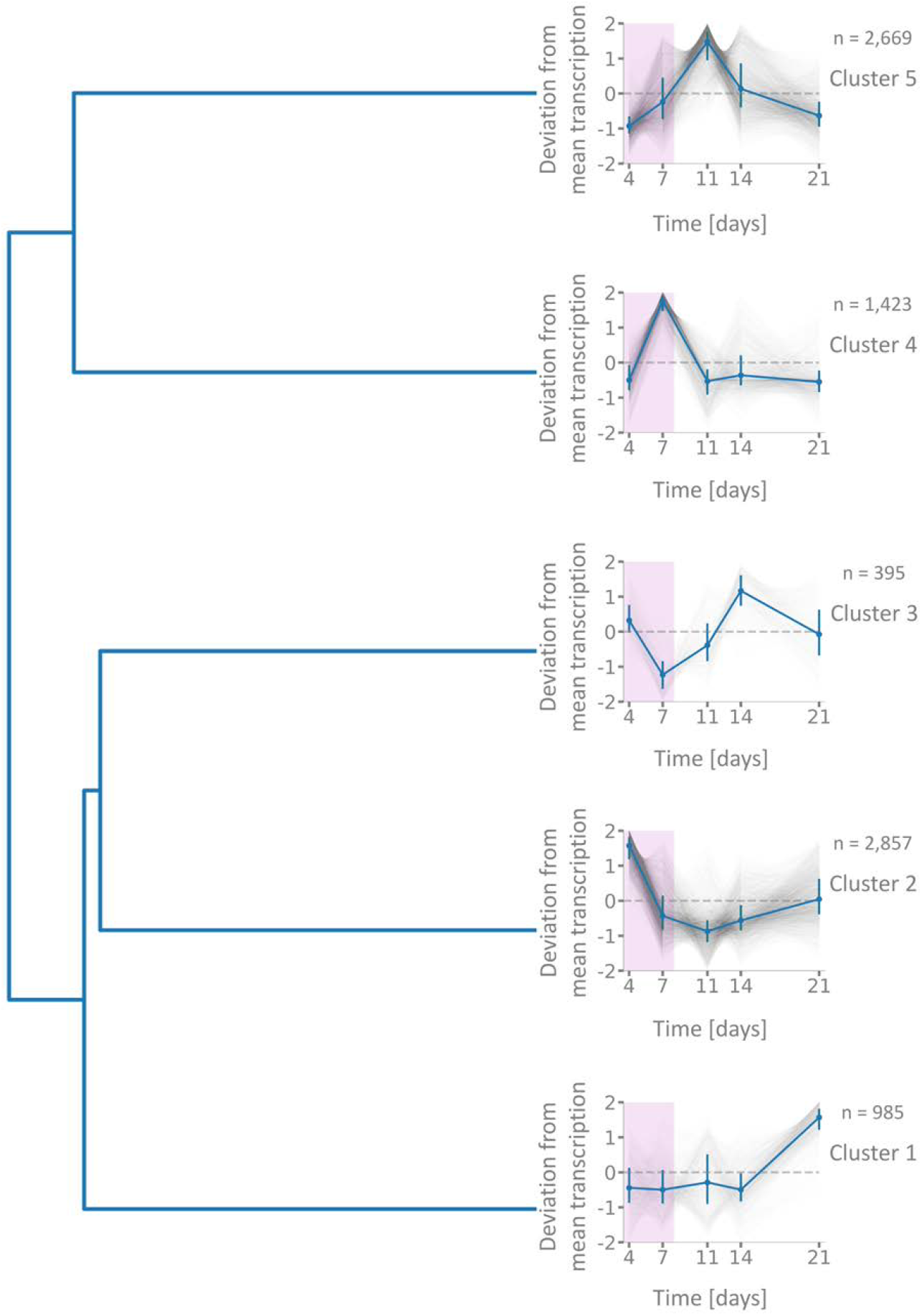
Temporal transcription patterns of 8,329 differentially expressed genes in the So ce836 genome. The deviation from mean transcription (z-scores) is plotted over growth time. A deviation value of 0 indicates that a gene is transcribed at its mean level calculated over all time points. A value of 2 indicates that transcription levels deviate from the mean by two standard deviations. The dendrogram shows the relationship between five transcription clusters based on hierarchical clustering with City Block distance and the average linkage method. The mean transcription of individual genes over time-point replicates is plotted in grey and median values for the whole transcription cluster are plotted in blue with error bars indicating first and third quartiles. The pink-shaded area indicates the exponential growth phase.

### Analysis of temporal transcription supported the identification of individual BGCs

Core biosynthetic genes within specific BGC regions commonly followed similar temporal transcription patterns (***Figure 3***, Suppl. Fig. 1), likely due to their organization in single, co-transcribed operons (Fischbach & Walsh, 2006). However, a few BGC regions predicted by antiSMASH exhibited deviating transcription of their core genes over time, including BGC regions 7, 20, 25, and 41 (Suppl. Fig. 2). Closer inspection showed that each of these regions likely constituted several BGCs (Suppl. Fig. 3), since their BGC core genes were spatially separated or located on different DNA strands. Consequently, we split BGC region 7 into two subregions (7_1 and 7_2), corresponding to antiSMASH-predicted core genes involved in the biosynthesis of resorcinol and a phenazine compound, respectively (Suppl. Table S1). Similarly, BGC region 20 was split into three subregions (20_1 - 20_3) and region 41 was split into two (41_1, 41_2). BGC subregion 41_1 was predicted to encode the biosynthesis of a redox-cofactor and the other subregions are likely involved in the production of NRPs or NRP/PK-hybrids (Suppl. Table S1). We assigned genes other than core biosynthetic genes within overlapping regions to both BGC subregions (Suppl. Fig. 3).

**Figure 3.**
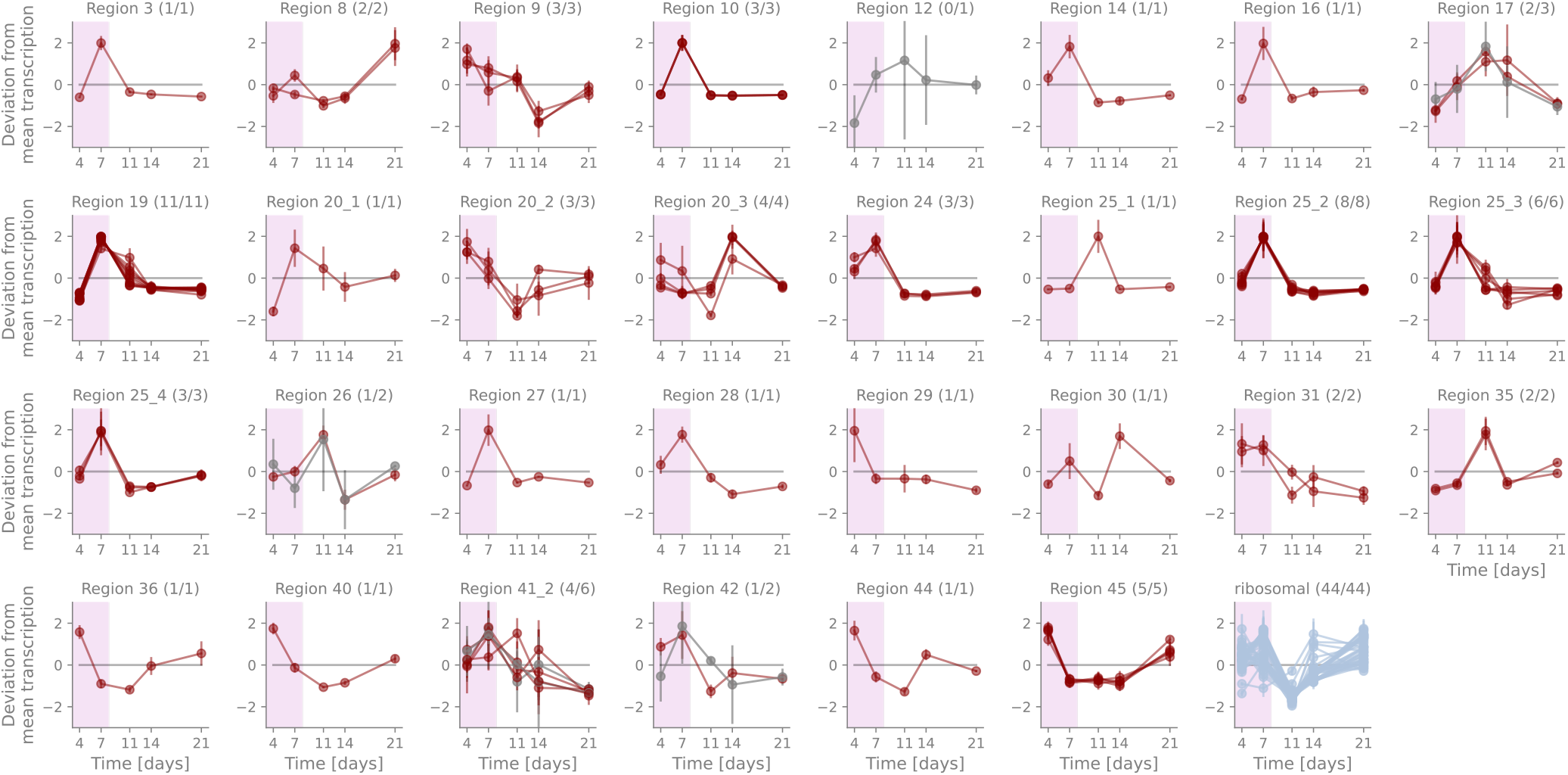
Temporal transcription of core biosynthetic genes of NRP- and PK-related BGCs. The deviation from mean transcription (z-score) is plotted over growth time for each BGC associated with the production of NRPs, PKs, or their hybrids. Numbers in brackets indicate the number of differentially expressed core genes and the total number of core genes in the BGC, respectively. Each curve represents the time-course of transcription of one core biosynthetic gene. Data points indicate the means over replicates at a time point and error bars represent the adjusted standard deviations. Differentially transcribed genes are represented in dark red, genes without a significant difference in transcription over the time-course experiment in grey. The last panel shows the temporal transcription of 44 genes encoding ribosomal proteins for comparison. Pink shading indicates the exponential growth phase.

Temporal transcription patterns did not always coincide with BGC boundaries, however. Region 25 included four spatially separated groups of BGC core genes (Suppl. Fig. 4), three of which encoded thioesterase at the C-terminal ends of their last core gene, suggesting independent biosynthetic pathways within this region (Schwarzer & Marahiel, 2001). Subregion 25_3 represents the previously described epothilone BGC with a 100% protein similarity to MIBiG entries (BGC0000989 - BGC0000991 (Julien et al., 2000; Molnár et al., 2000; Tang et al., 2000; L.-P. Zhu et al., 2013)). Moreover, we indeed detected epothilone A production by So ce836 via LC/MS analysis. Subregion 25_4 shares 62% similarity and an identical gene order with the glidopeptin BGC (MIBiG entry BGC0001608 (Wang et al., 2018)). While the separation of region 25 into several BGCs is therefore justified, temporal transcription patterns were largely identical for three of the four subregions (Suppl. Fig. 2).

### The transcription of most core genes for NRP and PK biosyntheses peaked during exponential growth

Among 307 genes from BGCs for NRP and PK production, 260 (85%) were differentially expressed over our time-course experiment. This is a significant over-representation compared to the number of differentially expressed genes among all genes (p-value = 1.41e-3). A particularly large proportion of core biosynthetic genes (93%, 75 out of 81 core genes) were differentially expressed (p-value = 3.34e-4).

The majority of differentially expressed core genes from BGCs for PK and NRP syntheses were maximally transcribed on day 7 (60%) or day 4 (20%) (***Figure 3, Figure 4***). Accordingly, 40 (53%) of the 75 differentially expressed core genes belonged to transcription cluster 4, and 18 (24%) to transcription cluster 2. This means a statistically significant overrepresentation of core genes among the two clusters, which both displayed a strong tendency for maximum transcription in the exponential growth phase (***Figure 2***, Suppl. Table S2). Transport-related genes and additional biosynthetic genes within BGCs often showed similar temporal trends of transcription as the core genes (Suppl. Fig. 5, Suppl. Fig. 6). In contrast, transcription of regulatory genes from these BGCs exhibited deviating temporal patterns (Suppl. Fig. 7). Of note, however, the smCOG-based prediction of these accessory genes is less reliable than the rule-based identification of core biosynthetic genes (Medema et al., 2011). Therefore, we focused on the latter for our further analyses.

**Figure 4.**
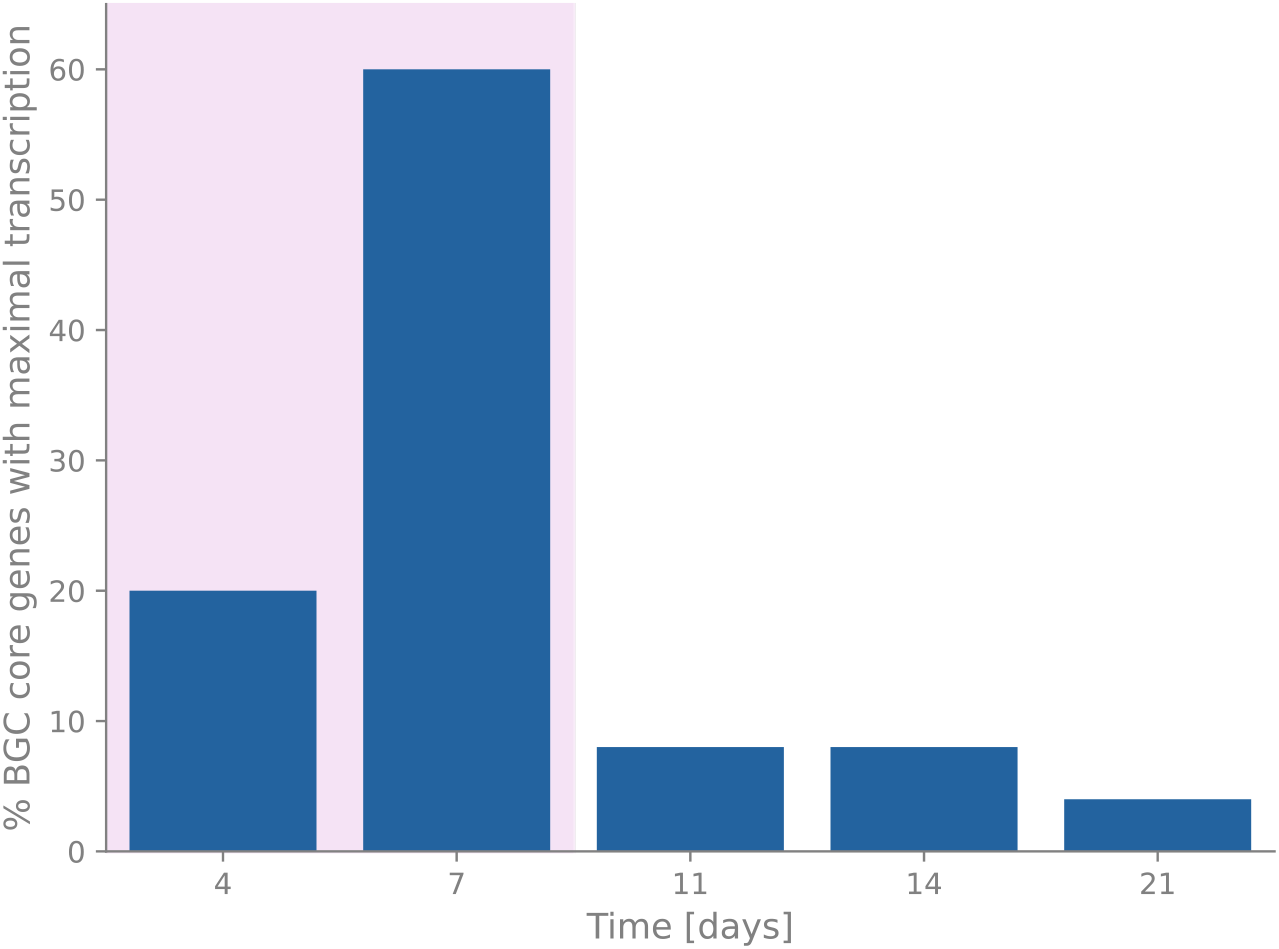
Temporal distribution of maximal gene transcription of BGC core genes associated with NRP- and PK-production. Out of 81 core genes from NRP- and PK-related BGCs, 75 were differentially expressed in our experiment. This figure shows for each time point how many of the differentially expressed core genes were maximally transcribed at that time. Pink shading indicates the exponential growth phase.

For comparison, we plotted the temporal transcription of fourty-four genes encoding ribosomal proteins (***Figure 3***, Suppl. Table S3). The transcription activity of most ribosomal protein genes also peaked during the exponential growth phase, but their temporal patterns nevertheless differed markedly from BGC core genes by showing additional transcription maxima on days 14 and 21 (***Figure 3***, Suppl. Fig. 8).

### Transcriptional patterns of other BGC classes

The So ce836 genome encodes 13 BGCs for potential RiPPs, six for terpenes, and one each for phosphonate, resorcinol, and phenazine compounds (Suppl. Table S1). The core genes of these BGC classes showed more diverse temporal transcription patterns compared to NRP- and PK-related BGCs (Suppl. Fig. 1, Suppl. Fig. 9, Suppl. Fig. 10). From the 21 core genes of RiPP-related BGCs, 15 were significantly differentially expressed and eight of these (53%) reached their maximal transcription level in the exponential phase (day 4 and day 7) (Suppl. Fig. 1). The BGCs associated with the production of terpenes contain six core genes (Suppl. Fig. 9), only two of which were maximally transcribed in the exponential phase, whereas two reached their maximal transcription level in the death phase (at day 21). The resorcinol- and phenazine-related BGCs are localized close to each other on the genome, but yet their temporal expression differed strongly, indicating different mechanisms of transcriptional regulation (Suppl. Fig. 10). So ce836 was also predicted to produce a phosphonate, encoded by BGC region 38. The corresponding core gene was most strongly expressed in the early exponential phase (day 4) and in the death phase (day 21).

### BGC core gene expression was correlated with metabolite net production rates

Strain So ce836 produced large amounts of the natural compound icumazole A. The concentration of icumazole A, as measured by HPLC based on its absorption of UV light (304 nm, 15.27 min retention time), was maximal on day 11 and decreased thereafter, suggesting the compound was catabolized by the bacterium or otherwise degraded (***Figure 5A***, blue curve). We determined the net compound production rate (arbitrary units) at each time point by assessing the slope of the concentration curve and then related it to the concentration of bacterial cells in the culture at the same time point (as shown in Figure 1) to estimate the net compound production rate per cell. As a result, we found that the net icumazole A production rate per myxobacterial cell peaked on day 7, i.e. during exponential growth (***Figure 5A***, green curve). Strikingly, this net cellular icumazole A production rate correlated strongly with the transcriptional pattern of the ten core genes from the icumazole BGC (r=0.88; (Xie et al., 2022)), since the maximum deviation from mean transcription was also reached on day 7 (***Figure 5A***, red curves).

**Figure 5.**
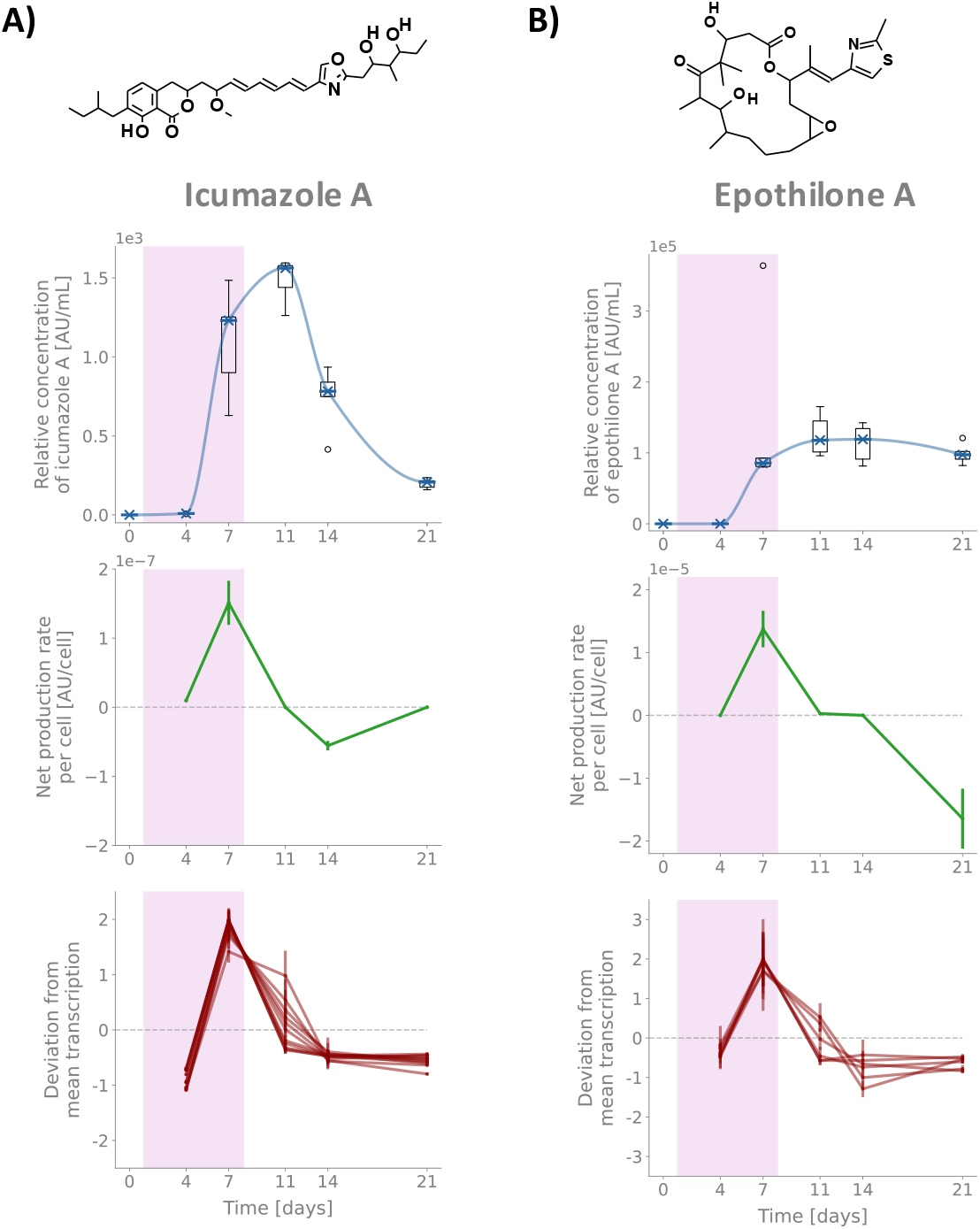
Net production rates of icumazole A (A) and epothilone A (B), and transcription of core genes in the underlying BGCs. The images above the plots show the chemical structures of icumazole A (left, PubChem CID 101557675 (National Center for Biotechnology Information, 2022b)) and epothilone A (PubChem CID 448799 (National Center for Biotechnology Information, 2022a)). The upper panels (blue) represent the relative concentration of natural products estimated from LC/MS analyses. Boxplots over five replicates per time point are shown. The curves were obtained by monotone piecewise cubic interpolation over median concentrations. The second panels (green) show the net compound production rate per myxobacterial cell, based on the slope of the compound concentration per mL and the mean cell count per mL at a time point. Error bars indicate propagated errors. The lower panels (red) represent the deviation from mean transcription (z-scores) of core biosynthetic genes in the icumazole and epothilone BGCs, respectively. The zero-line marks the mean transcription of a gene over all time points. Pink shading indicates the exponential growth phase.

At the same time, we also detected epothilones in the metabolome of So ce836. Epothilone A was only weakly detectable by UV absorption (250 nm, 9.05 min retention time) at most time points, and since quantification of epothilone A by UV absorption and mass spectroscopy gave similar results for days 4-14 (Suppl. Fig. 11), we relied on LC/MS mass peaks (494.2571 m/z, 9.05 min retention time) to estimate the relative concentration of epothilone A. Similar to icumazole A, the concentration of epothilone A in the growth medium was highest in the stationary phase (***Figure 5B***, blue curve). Again, the net cellular production rate of epothilone A was maximal on day 7, and it correlated with the time course of transcription of epothilone BGC core genes (r=0.78) (***Figure 5B***, green and red curves).

Taken together, we observed that bursts of BGC core gene transcription were accompanied by distinct surges in net production rates per cell for the two known compounds icumazole A and epothilone A. Even though temporal dynamics of normalized BGC core gene transcription and compound production rates were highly similar for these compounds (***Figure 5***), absolute transcription levels of the core genes (i.e. sequencing read counts per kilobase) differed more than 100-fold between associated BGCs (Suppl. Table S1, Suppl. Fig. 8). Notably at maximum transcription on day 7, the median number of normalized sequencing read counts mapped to the icumazole BGC core genes was 427 per kilobase (1,732 to 12,798 reads per gene), and for the epothilone BGC was only 3 per kilobase (13 to 55 reads per gene) (***Figure 6***, Suppl. Fig. 8). And on day 4, when almost no icumazole A net production was detectable, the median number of sequencing reads mapped to icumazole BGC core genes was at its minimum of 15 per kilobase (55 to 399 reads per gene), but yet this number was larger than the maximal transcription level of the epothilone BGC core genes (Suppl. Fig. 8). The transcript levels of other BGCs were intermediate between those of epothilone and icumazole BGCs, except for four BGCs with slightly lower read counts than the epothilone BGC and five BGCs that had even higher read counts per kilobase than the icumazole BGC, although we were unable to detect their (as yet unknown) natural products (***Figure 6***, Suppl. Table S1).

**Figure 6.**
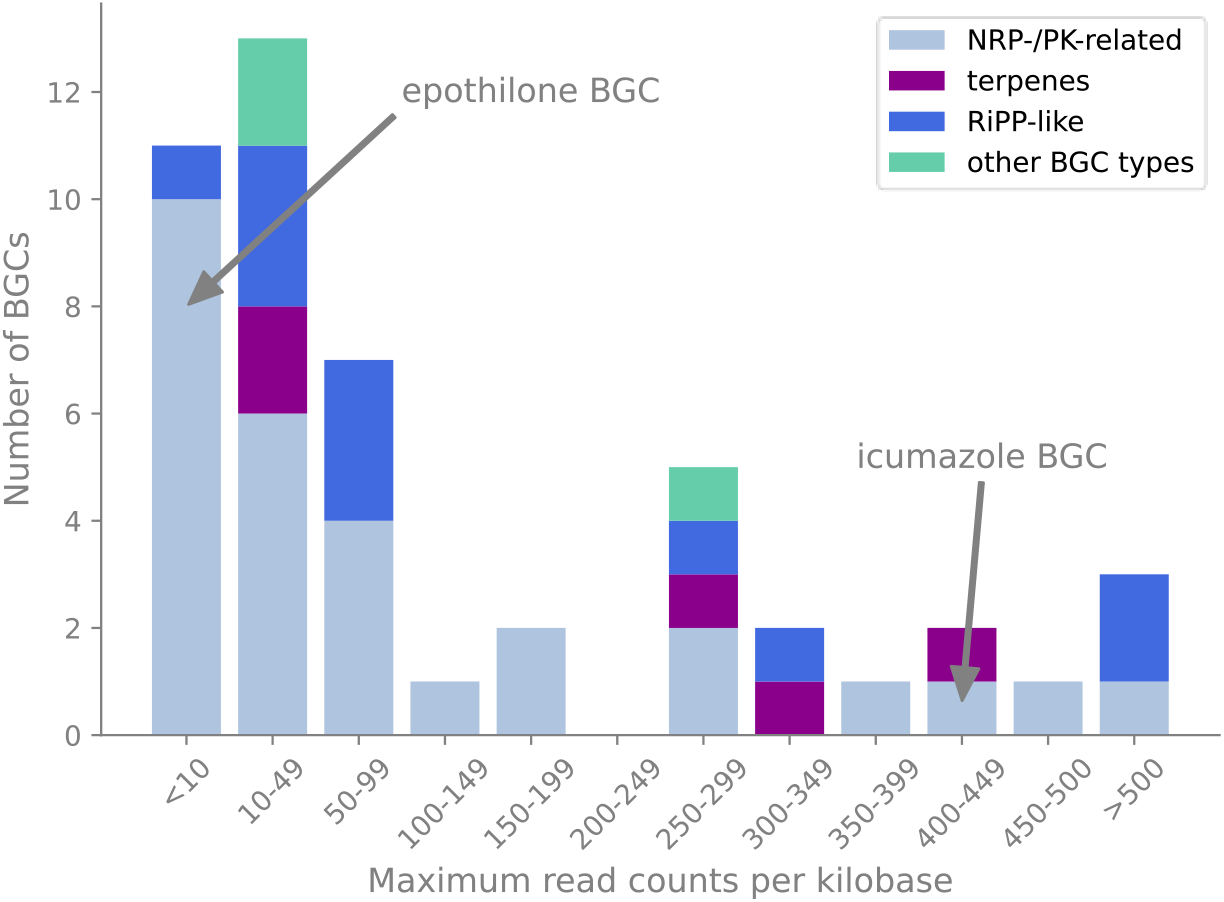
Distribution of BGC read counts on their day of maximal transcription. For each BGC core gene the mean read counts per kilobase after DESeq2 normalization was calculated over all replicates on the day of maximal BGC transcription. Plotted here is the median over all core genes in a BGC. Each type of BGC is represented in a different color. The ‘other BGC types’ include resorcinol-, phenazine-, and phosphonate-producing BGCs. Please note that the first two bins and the last one do not have the same range as the other bins. With a maximal read count per kilobase of 3, the epothilone BGC is located in the left most bin. The icumazole BGC has a maximum read count per kilobase of 427.

While the net cellular production rates of both icumazole A and epothilone A were maximal during exponential growth, the highest relative concentrations of the compounds were reached in later growth phases (***Figure 5***). This might be due to the accumulation of compounds over time and the large, metabolite-producing biomass available in the batch culture towards the end of the growth phase.

### Functional analysis of transcription clusters based on COG categories

To assess the temporal transcription in a genome-wide context with respect to cellular function, we classified the proteins encoded in the So ce836 genome into Clusters of Orthologous Groups (COG; Suppl. Table S4) (Galperin et al., 2015). Out of 8,329 differentially expressed genes, we could associate 6,081 (73%) to COG categories using the eggNOG-mapper (Cantalapiedra et al., 2021). We then performed an over-/under-representation analysis of COG categories within each of the five transcription clusters (Suppl. Table S5). Among genes in transcription cluster 2 (maximally transcribed on day 4, i.e. during early exponential growth), COG categories *M* (‘Cell wall/membrane/envelope biogenesis’) and *F* (‘Nucleotide transport and metabolism’) were significantly over-represented compared to their distribution among all annotated and differentially expressed genes in So ce836 (for adjusted p-values, see Suppl. Table S5). In transcription cluster 4 (maximally transcribed on day 7), categories *P* (‘Inorganic ion transport and metabolism’), *J* (‘Translation, ribosomal structure and biogenesis’), and *Q* (‘Secondary metabolites biosynthesis, transport and catabolism’) were significantly over-represented. In contrast, in transcription cluster 5 (maximally transcribed on day 11, i.e. during the early stationary phase), COG categories *J, M F, P*, *E* (‘Amino acid transport and metabolism’), and *H* (‘Coenzyme transport and metabolism’) were significantly under-represented, while category *O* (‘Posttranslational modification, protein turnover, chaperones’) was over-represented. Hence, the distribution of genes classified to COG category *Q* (‘Secondary metabolites biosynthesis, transport and catabolism’) confirmed our previous notion of BGC transcription mostly during exponential bacterial growth. In transcription cluster 3 (maximally transcribed on day 14), no COG category was significantly over- or under-represented, and in cluster 1 (maximally transcribed on day 21), category *G* (‘Carbohydrate transport and metabolism’) was over-represented.

## Discussion

Our data show that the transcription of BGCs in *Sorangium* sp. is determined by bacterial growth. In a batch culture, the transcription level of most BGC core genes (PKSs and NRPSs in particular) was maximal during exponential growth, when the cellular metabolism was simultaneously engaged in the synthesis of cell wall components, proteins and nucleic acids, and in trans-membrane transport activities (as indicated by the functional analysis of genome-wide transcription patterns). As soon as bacterial growth ceased and the culture reached the stationary phase, BGC transcription quickly levelled off. Strikingly, transcriptional bursting of BGC core genes appeared to directly determine the net synthesis rates of the NRP/PK-hybrids epothilone A and icumazole A. Such a correlation between natural compound synthesis and BGC transcription explains the success of previous attempts to boost compound production by increasing the level of transcription (Peng et al., 2018; Rachid et al., 2007, 2009; Ye et al., 2019). This result also suggests that biosynthesis proteins may get degraded (Trötschel et al., 2013) or their activities downregulated during later growth stages, which coincides with the observation of increased transcriptional activities of genes involved with protein modification and turnover in the early stationary phase. The close association between BGC transcription and compound production observed here for a wild-type *Sorangium* strain warrants efforts towards developing improved genetic engineering tools for this organism, since the molecular manipulation of transcriptional regulation may be a viable strategy for improving compound yields.

Of note, absolute transcription levels varied widely between BGCs, including those for epothilone and icumazole. It is currently unknown which level of BGC transcription may be required for natural compound production. Our results indicate that the transcription level may vary over several orders of magnitude among active BGCs, even within a single genome, suggesting it may be difficult to find any universal threshold for transcription above which the production of detectable amounts of natural compounds could reliably be predicted (Mungan et al., 2022). Our data also show that BGC read counts at single time points may provide limited information about compound synthesis. Rather, time course data were required to resolve the transcriptional bursts that were accompanied by increased rates of compound production in *Sorangium* sp.

One limitation of our analysis is that it relied on measurements of net production rates of compounds from the bulk of the bacterial culture, whereas we could not directly assess the underlying synthesis rates within bacterial cells. Decay of icumazole A clearly dominated over its synthesis during the late stages of the batch culture, but it is unknown whether the compound was equally unstable while the bacteria were growing or whether it was actively degraded. Hence, the measured net production rates likely resulted from both synthesis and concomitant degradation, adding complexity to understanding their genetic regulation. Arguably however, the net production rate is the most relevant measure for biotechnology.

Core biosynthesis genes from 48 BGCs (out of 52 BGCs in total) in *Sorangium* sp. So ce836 were differentially expressed over our time-course experiment, i.e. their transcription levels varied significantly over time. Moreover, this was true for all core genes of the majority of BGCs, and their temporal transcription patterns were largely synchronous within BGCs. These results indicate that the majority of BGCs in the genome were transcriptionally active, even though we had applied only a single cultivation condition. Similarly, an earlier investigation applying microarray and proteome analyses had demonstrated that the majority of BGCs in an *M. xanthus* culture were expressed (Bode et al., 2009; Schley et al., 2006), and hence, this may be a common pattern in myxobacteria. Note, however, that our present study provides rich insights into genome-wide temporal patterns of gene activity that were not resolved previously on the basis of microarray hybridization or quantitative PCR (Bode et al., 2009). Generally, the temporal dynamics of BGC transcription, translation, and natural compound assembly in bacteria are poorly understood (Machado et al., 2017). Following the transcription of BGC genes, the synthesis of natural compounds may be regulated at the translational or posttranslational levels, or limited by the supply of building blocks, cofactors, or energy (Wenzel & Müller, 2009).

It must also be taken into consideration that an observed discrepancy between the genomic endowment with biosynthetic genes and the yield of natural compounds may be caused by limitations of the techniques for metabolite extraction and analysis (Amos et al., 2017; Panter et al., 2021). Due to the inherent complexity of metabolome data, it is generally challenging to discover unknown natural compounds and associate them with their BGCs (Panter et al., 2021). However, if the net cellular production of natural compounds is correlated with BGC transcription also in other cases similar to epothilone and icumazole, we suggest that temporal transcriptome patterns may provide useful criteria for the filtration of time-resolved metabolome data.

Our observation of high natural compound production rates in growing myxobacterial cells challenges the frequently cited notion of preferred BGC expression during late bacterial growth phases, which is based on observations on *Streptomyces* spp. and other actinobacteria (van der Heul et al., 2018; van Wezel & McDowall, 2011; Wohlleben et al., 2017). It suggests that adjusted batch runtimes, product retention approaches, or even continuous fermentation might offer the potential for maximization of biotechnological product yields from myxobacteria (Frykman et al., 2005).

To our knowledge, our study for the first time relates the temporal dynamics of BGC expression to patterns of genome-wide transcription in any myxobacterium. A more detailed understanding of the diversity of BGC expression and regulation will facilitate the discovery of novel natural compounds, the induction of their synthesis, and the optimization of their large-scale production.

## Methods

### Liquid cultures of So ce836

Pre-cultures of *Sorangium* sp. So ce836 were grown in CYH-medium (Hoffmann et al., 2018) for 4 days at 28°C with shaking at 160 rpm. Subsequently, 90 mL H-medium (Hoffmann et al., 2018) were inoculated with 10 mL of pre-culture, with five replicates for each growth period (0, 4, 7, 11, 14, and 21 days). Culture flasks were incubated at 28°C with shaking until the corresponding growth time was reached. The cultures were then homogenized with a MICCRA MiniBatch D-9 homogenizer (full power, 30-40 sec), equipped with a MICCRA PICO DS-8-P dispersing tool (MICCRA GmbH).

### Cell counting

100 μL homogenized culture was mixed with 900 μL 4% formaldehyde (Sigma-Aldrich) and stored at 4°C. 50 μL of the formaldehyde-fixed cells were added to 1 mL PBS (Carl Roth) and 450 μL methanol and incubated for 15 min in an ultrasonic bath at 35°C. 500 μL of this solution was added to 10 mL sterile filtered PBS, stained with 2 μL SYBR Green (Lonza), incubated for 10 min at room temperature, and filtered on a black polycarbonate filter (Merck Millipore). 25 μL 0.25% DABCO-glycerol-solution (Sigma-Aldrich) was added, the filter was covered with covering glass, and the cells counted under a fluorescent microscope (Zeiss, 1000x, GFP-filter). For each sample (six time points with five biological replicates each), cells were counted in ten microscopic fields (technical replicates).

### RNA extraction

Immediately after removal of 100 μL homogenized culture for cell counting, the cells in the remainder of the culture were pelleted by centrifugation for 5 min at 11,000 rpm. The supernatant was retained for metabolite extraction (see below). Four to five inoculation loops of cells were taken from the pellet, added to 2 mL RNA*later* (Ambion) solution, and stored at 4°C. The remaining pellet was remixed with the supernatant for metabolite extraction (see below). RNA was extracted with the RNeasy Mini kit (Qiagen). After removal of residual DNA with DNase *Turbo* (Ambion), RNA was precipitated with three volumes of 100% ethanol at −20°C, redissolved in RNase-free water and quantified by using the *Quant-iT RiboGreen* RNA assay (Fisher Scientific) and a microplate fluorescence reader (Tecan). Ribosomal RNA was depleted by applying the *Ribo-off* kit (Absource) and the quality and quantity of resulting RNA was assessed following electrophoresis on a *Bioanalyzer 2100* instrument equipped with a *6000 Pico Chip* (Agilent).

### Metabolite extraction

XAD 16 (Amberlite) was washed with H_2_O, acetone, and methanol to remove polymers and was then stored in H_2_O. For each culture, a gauze bag with 2% (v/v) of XAD 16 was made and added to the supernatant and the rest of the cell pellet. After incubation at 4°C overnight, the gauze bag was removed and rinsed with deionized water. The cells were pelleted by centrifugation of 5 min at 11,000 rpm and the supernatant removed. Cell pellet and the XAD bag were incubated together for 1 hour in 100 mL acetone. Cells and XAD were removed by filtration through paper, the acetone solution collected, and subsequently vaporized with a rotary evaporator (Heidolph Laborota 4003 control, heat sink at 15°C, water bath at 40°C). The remaining residue was dissolved in 1 mL methanol, the solution was centrifuged for 10 min at 11,000 rpm, and the supernatant was used for LC/MS analyses.

### LC/MS analysis

For HPLC/MS analyses, 50 μL of methanol supernatant (see above) was mixed with 50 μL pure methanol. The HPLC measurements were performed using an Agilent 1260 series with DAD detector (Agilent Technologies) and a XBridge BEH C18, 130 Å, 3.5 μm, 2.1 mm x 100 mm column (Waters). For compound separation we used a linear gradient from (A) H_2_O + 5% acetonitrile + 5mM ammonium acetate + 0.004% acetic acid to (B) acetonitrile + 5% H_2_O + 5 mM ammonium acetate 0.004% acetic acid at a flow rate of 0.3 mL/min at 40°C. The gradient was initiated with 90% A and 10% B, increasing to 100% B within 30 min, followed by 100% B for 10 min, and a reequilibration to 90% A and 10% B. UV spectra were created with a range of 200 - 600 nm. Mass spectra were obtained with a maXis UHR-TOF-MS (Bruker Daltonics) on an ACQUITY UPLC BEH C18, 130Å, 1.7 μm, 2.1 mm X 50 mm column (Waters) with an ACQUITY UPLC BEH C18 VanGuard, 130Å, 1.7 μm, 2.1 mm X 5 mm pre-column (Waters). The gradient was set from (A) 0.1% formic acid water solution to (B) 0.1% formic acid acetonitrile solution at a flow rate of 0.6 mL/min. The gradient was established with 95% A and 5% B for 0.5 min, increasing solvent B to 100% within 19.5 min, and followed by 100% B for 10 min at a temperature of 40°C.

We identified compounds by comparison of molecular weight, retention time, and UV spectrum (where available) to known natural compounds from an in-house database. In order to estimate the relative concentration of product per mL sample we used either the area under the UV peak (where available) or the mass peak at the corresponding retention time and adjusted it by the dilution factor from sample preparation.

For calculating the net cellular production rate for icumazole A and epothilone A, we first used monotone piecewise cubic interpolation (scipy.interpolate.pchip) on the median concentrations of product per mL (arbitrary units/mL) over time. The derivative at each time point (i.e. the slope of the interpolation curve) indicated the net production rate per day. Through division by the concentration of bacterial cells, we obtained the net cellular production rate.

### Library preparation and RNA sequencing

Libraries for RNA sequencing were prepared by applying the TruSeq Stranded mRNA kit (Illumina) according to the manufacturer’s protocol, and sequenced on a NextSeq 550 instrument (Illumina) by using mid-output flow cells with 300 cycles.

### Genome sequencing

A complete genome sequence was generated combining SMRT long-read sequencing (Pacific Biosciences) and Illumina short-read sequencing (Illumina) as described in detail previously (Steglich et al., 2018).

### Genome assembly, annotation, and BGC prediction

From PacBio *RSII* long-read sequencing the genome of So ce836 (GenBank accession: CP102233) was assembled using the Hierarchical Genome Assembly Process 3 Protocol (HGAP3, (Chin et al., 2013)) included in SMRTPortal 2.3.0. Illumina short-read sequences were preprocessed with fastp (version 0.20.1, (Chen et al., 2018)). Error-correction was performed by a mapping of Illumina short reads onto the finished genome using Burrows-Wheeler Alignment bwa 0.6.2 in paired-end (sample) mode and default settings (H. Li & Durbin, 2009) with subsequent variant and consensus calling using VarScan 2.3.6 (Koboldt et al., 2012). The final genome annotation was achieved with Prokka 1.14.6 (Seemann, 2014). For the prediction of BGC regions, we used antiSMASH version 6.1.1 (Blin et al., 2021).

### Differential gene expression analysis

The raw reads were preprocessed with fastp (version 0.20.1, (Chen et al., 2018)) and mapped to the reference genome (GenBank accession: CP102233) using an in-house script based on BWA-MEM (version 0.7.12, (H. Li, 2013)) and SAMtools (version 0.1.19, (Danecek et al., 2021)). The number of reads mapping to a gene were counted with HTSeq-count with mode union and stranded reverse option (version 0.13.5, (Anders et al., 2015)). For the analysis of genes differentially expressed over the time-course experiment, we implemented an in-house R script based on DESeq2 (version 1.32.0, (Love et al., 2014)). For further analyses, we used the normalized read counts generated by DESeq2. The independent filtering method of DESeq2 filtered out 610 of the 10,707 genes (6%) due to low read counts. For the remaining 10,097 genes we used DESeq2’s likelihood ratio test to identify the significantly differentially expressed genes.

The data discussed in this publication have been deposited in NCBI’s Gene Expression Omnibus (Edgar, 2002) and are accessible through GEO Series accession number GSE217497 (https://www.ncbi.nlm.nih.gov/geo/query/acc.cgi?acc=GSE217497).

In order to compare transcription patterns of genes with different absolute transcription levels, we used the z-score normalization (in the script referred to as deviation from mean transcription) on the mean over the biological replicates of each gene. In short, the z-score indicates the standard deviation from the mean transcription.

We implemented an in-house python script based on the python library scipy.cluster.hierarchy with the City Block metric and average linkage method, to cluster the differentially expressed genes by their deviation from mean transcription (z-scores) over the growth times.

To calculate the significance of BGC core genes among the transcription clusters we performed an over-/under-representation analysis based on the hypergeometric function. As the background distributions we used the number of genes within a transcription cluster over all differentially expressed genes.

### Functional analysis

The Clusters of Orthologous Groups of proteins (COGs) database (Galperin et al., 2015) provides a classification of proteins by their function. On its highest level, it groups proteins into 26 different categories (Suppl. Table S4). We obtained the COG categories for all predicted genes in So ce836 with the eggNOG-mapper (version 2.1.9, (Cantalapiedra et al., 2021; Huerta-Cepas et al., 2019)). We performed an over-/under-representation analysis to calculate the significance of the distribution of COG categories over transcription clusters. For this, we implemented an in-house python script based on the hypergeometric function with the distribution of COG categories over all differentially expressed genes with COG annotations as background distributions.

## Acknowledgements

This work was partially funded by the German Center for Infection Research (DZIF).

The authors thank Cathrin Spröer for genome sequencing and Nicole Heyer and Simone Severitt for excellent technical assistance.

