## supplement for "Bursts in biosynthetic gene cluster transcription are accompanied by surges of natural compound production in the myxobacterium *Sorangium* sp"

### Supplementary material

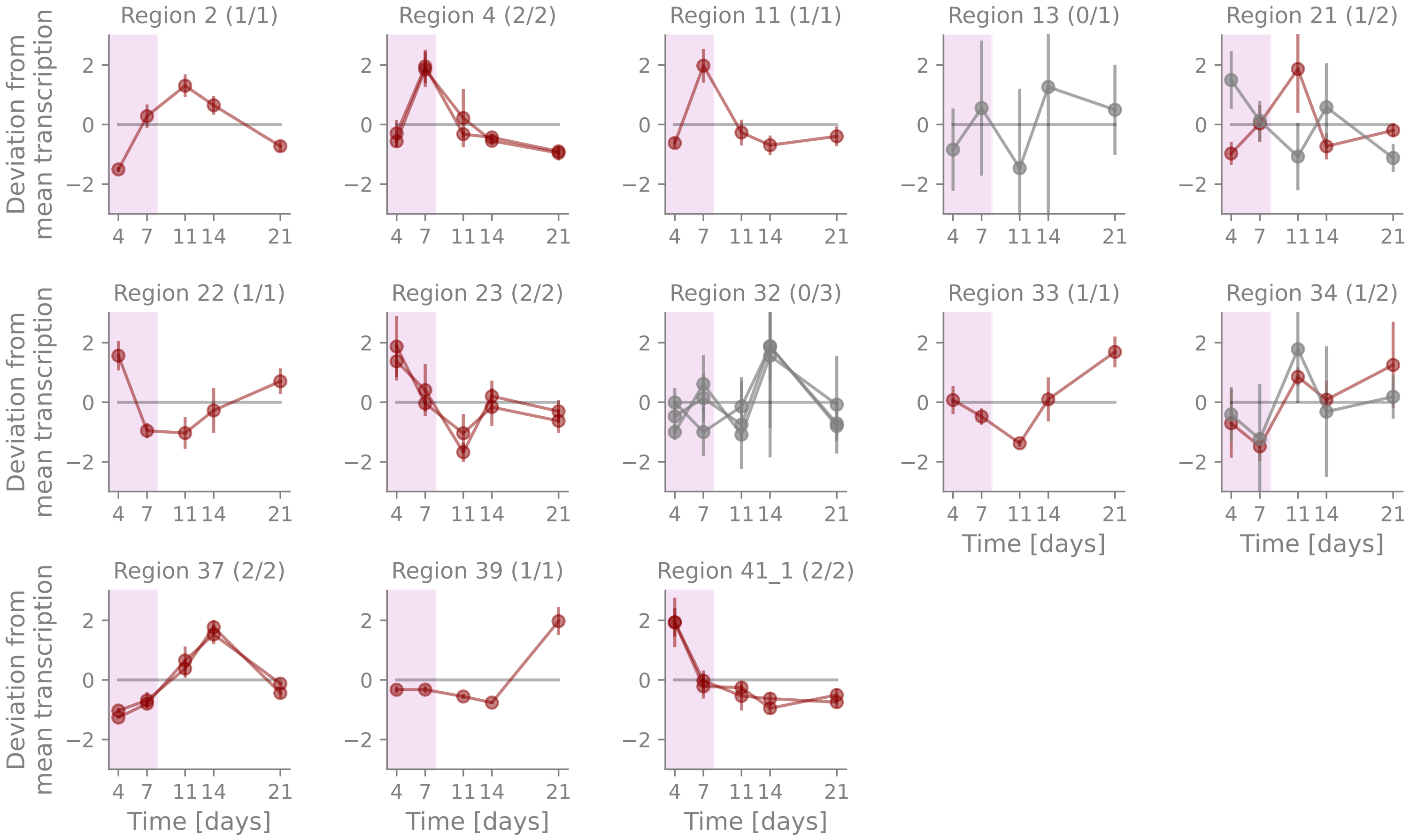


*Suppl. Fig. 1.* Temporal transcription of core genes of RiPP-related BGCs. Each subplot corresponds to one of the 13 antiSMASH-predicted BGC regions associated with the production of RiPPs (including the antiSMASH types RiPP-like, RRE-containing, thioamitides, LAP, microviridin, redox-cofactor). The antiSMASH-predicted BGC region 41 contains several BGCs and was split in two subregions. Subregion 41_1 corresponds to a potential redox-cofactor producing BGC. The numbers in brackets show the number of differentially transcribed core genes and the total number of core genes in the BGC (sub)region. The x-axis represents the growth time in days, the y-axis the deviation from mean transcription (z-score). Each curve corresponds to one core gene. The data points show the mean over the replicates at the time point, the error bars the adjusted standard deviation. Differentially transcribed genes are represented in dark red, genes without a significant difference in transcription over the time-course experiment in grey. The pink shading indicates the exponential growth phase.

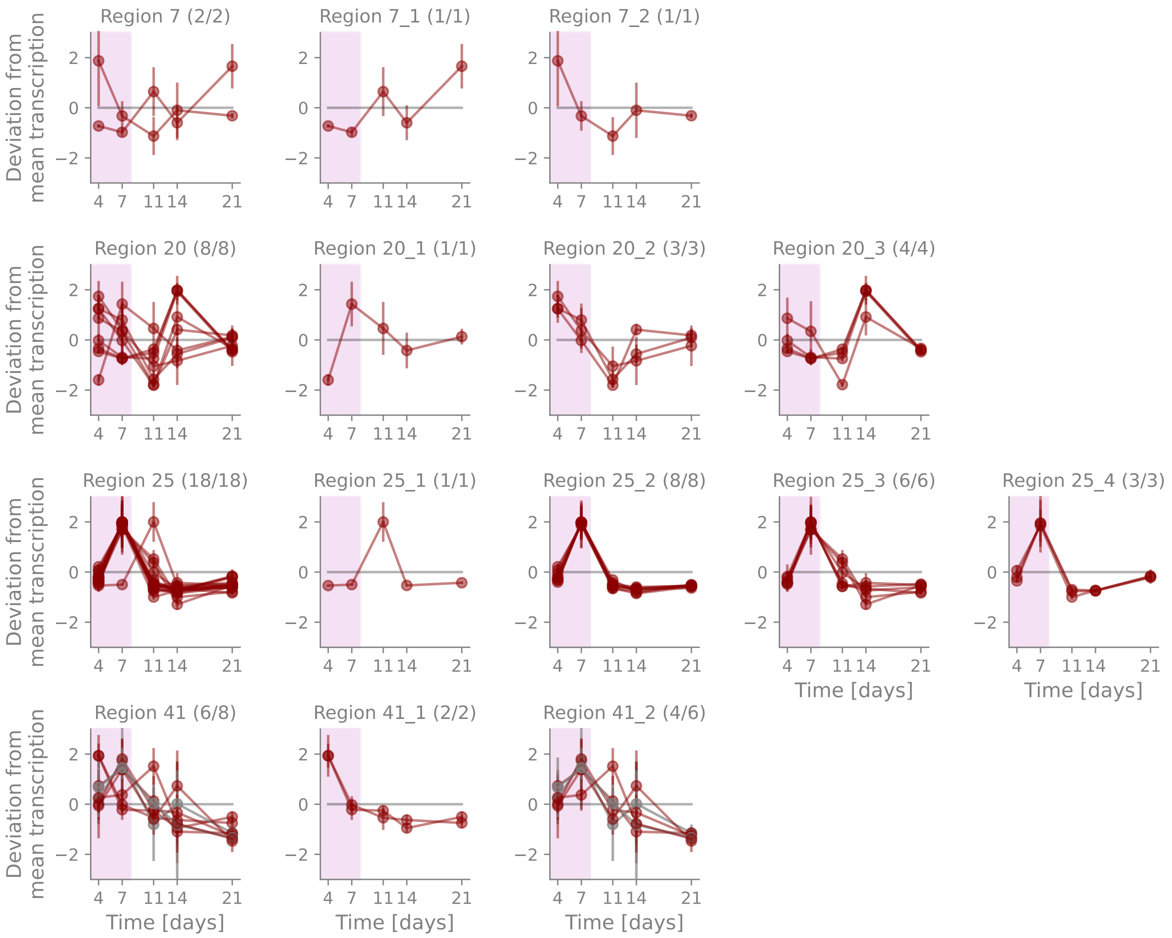


*Suppl. Fig. 2*. Temporal transcription of core genes of BGC regions with diverse transcription patterns. Each row corresponds to a different antiSMASH-predicted BGC region. The left most subplot shows the temporal transcription of all core genes in the region. The following subplots in each row display the core genes of a postulated BGC within the BGC region. The numbers in brackets show the number of differentially transcribed core genes and the total number of core genes in the BGC (sub)region. The x-axis represents the growth time in days, the y-axis the deviation from mean transcription (z-score). Each curve corresponds to one core gene. The data points show the mean over the replicates at the time point, the error bars the adjusted standard deviation, and the pink shading the exponential growth phase. Differentially transcribed genes are represented in red, genes without a significant difference in transcription over the time-course experiment in grey.

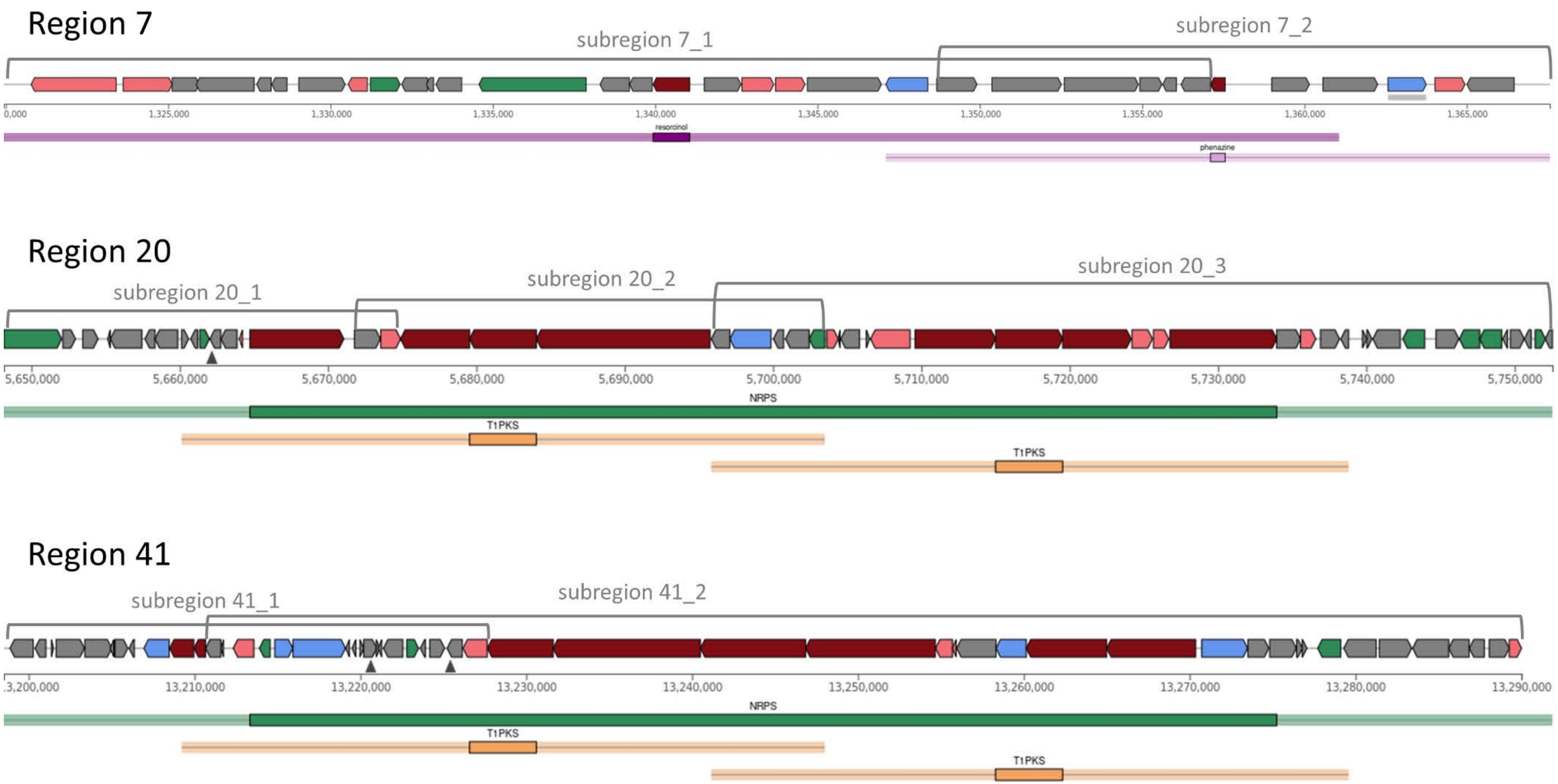


*Suppl. Fig. 3.* Structure of the antiSMASH-predicted BGC regions with diverse temporal transcription patterns of their core genes. Each arrow represents a gene. The direction of the arrow indicates the DNA strand the gene is located on. Core biosynthetic genes are shown in dark red, additional biosynthetic genes in coral, regulatory genes in green, transport-related genes in blue, and other genes in grey. The bars underneath the structure of the region marks the different types of BGCs identified by antiSMASH (resorcinol in purple, phenazine in pink, NRPS in green, type 1 PKS (T1PKS) in yellow). The grey brackets show the borders of the estimated BGCs within the BGC regions.

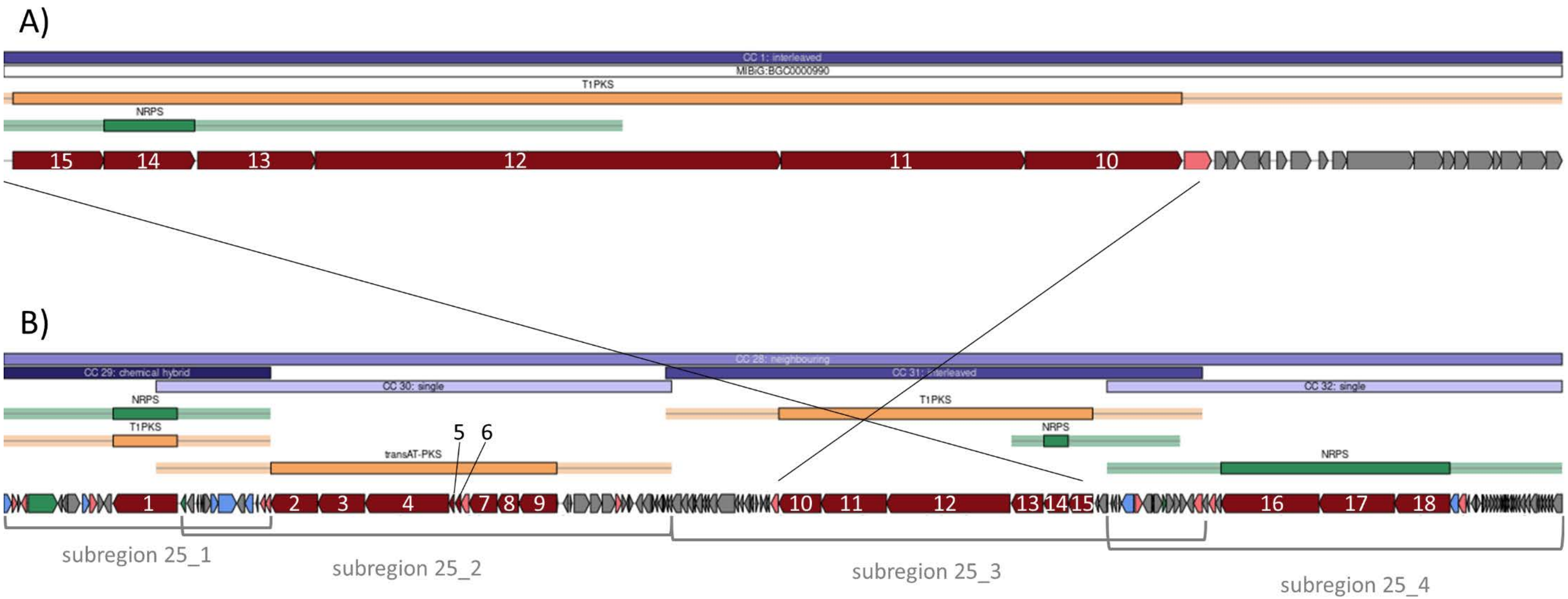


*Suppl. Fig. 4.* MIBiG epothilone BGC and antiSMASH-predicted BGC region 25 from So ce836. The bars in light and dark blue on the top of each plot indicate the candidate clusters predicted by antiSMASH. The yellow bars represent PKS-associated clusters (here T1PKS or transAT-PKS). The green bars correspond to NRPS-associated clusters. The bar underneath shows the genes in the cluster according to their location on the genome. Each arrow represents a gene. The direction of the arrow indicates the DNA-strand the gene is located on. Core biosynthetic genes are represented in dark red, additional biosynthetic genes in coral, transport-related genes in blue, regulatory genes in green, and all other genes in grey. **A)** One of the epothilone BGCs in MIBiG (BGC0000990) that corresponds to part of the antiSMASH-predicted region 25 in So ce836. **B)** The antiSMASH-predicted region 25 of So ce836. The eighteen core genes are numbered (1-18). The core genes 10 to 15, as well as the additional biosynthetic gene next to core gene 10 are orthologues of the core genes and the additional biosynthetic gene in the MIBiG epothilone BGC. The grey bars underneath the representation of the whole BGC region shows the ranges of the four BGC subregions (25_1 - 25_4) we proposed.

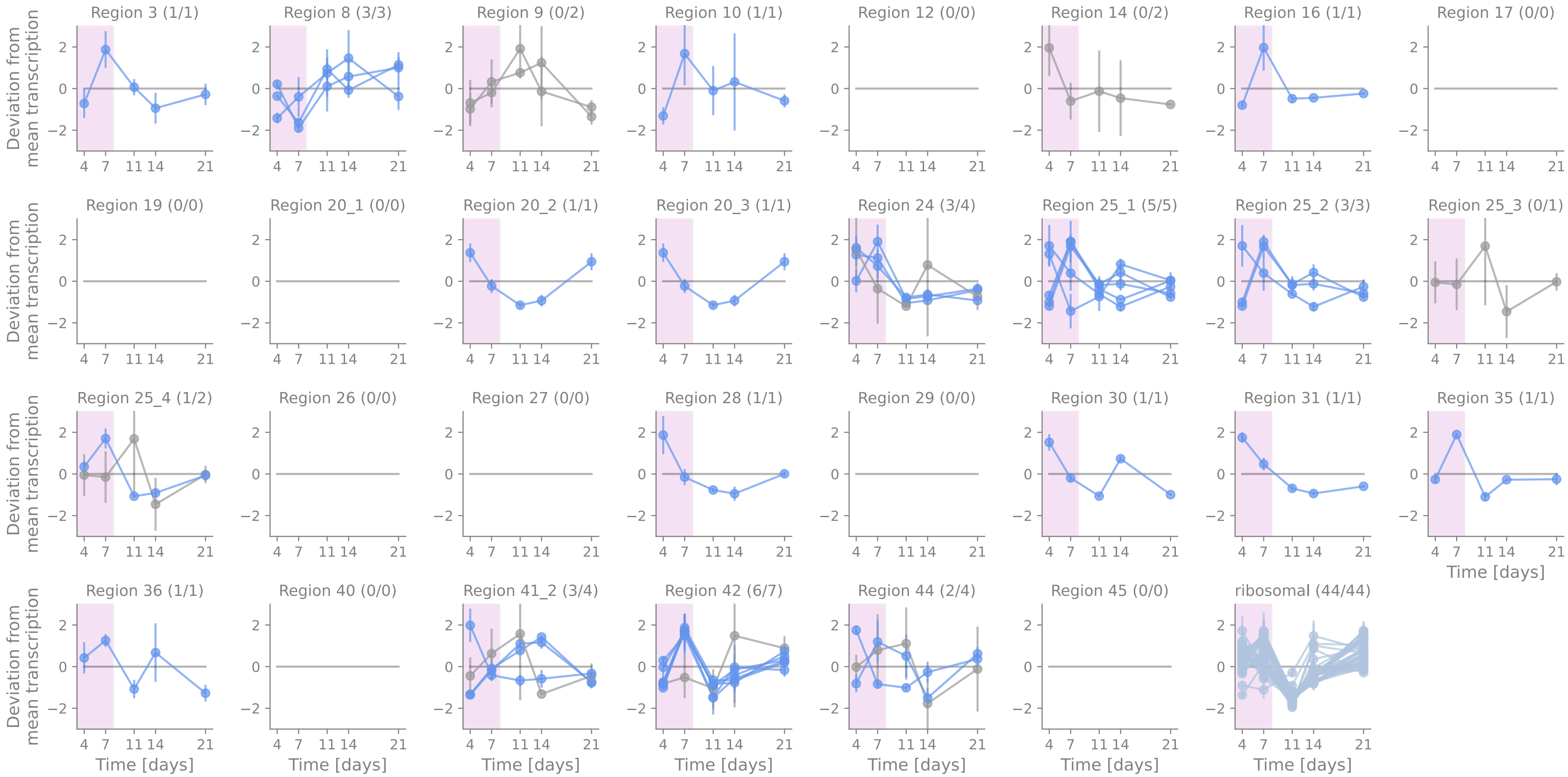


*Suppl. Fig. 5.* Temporal transcription of transport-related genes of NRP-, PK-related BGCs. Each subplot corresponds to one of the 25 antiSMASH-predicted BGC regions associated with the production of NRPs, PKs, or their hybrids. Some BGC regions contain several BGCs. We split these into the potential BGC subregions and represent each as an individual subplot indicated with underscores and subregion numbers. The numbers in brackets show the number of differentially transcribed transport-related genes and the total number of transport-related genes in the BGC (sub)region. The x-axis represents the growth time in days, the y-axis the normalized gene transcription level (z-score). Each curve corresponds to one transport-related gene. The data points show the mean over the replicates at the time point, the error bars the adjusted standard deviation, and the pink shading the exponential growth phase. Differentially transcribed genes are represented in blue, genes without a significant difference in transcription over the time-course experiment on grey. Genes filtered out by DESeq2 independet filtering are not represented. The last subplot shows the temporal transcription of 44 ribosomal genes as a comparison.

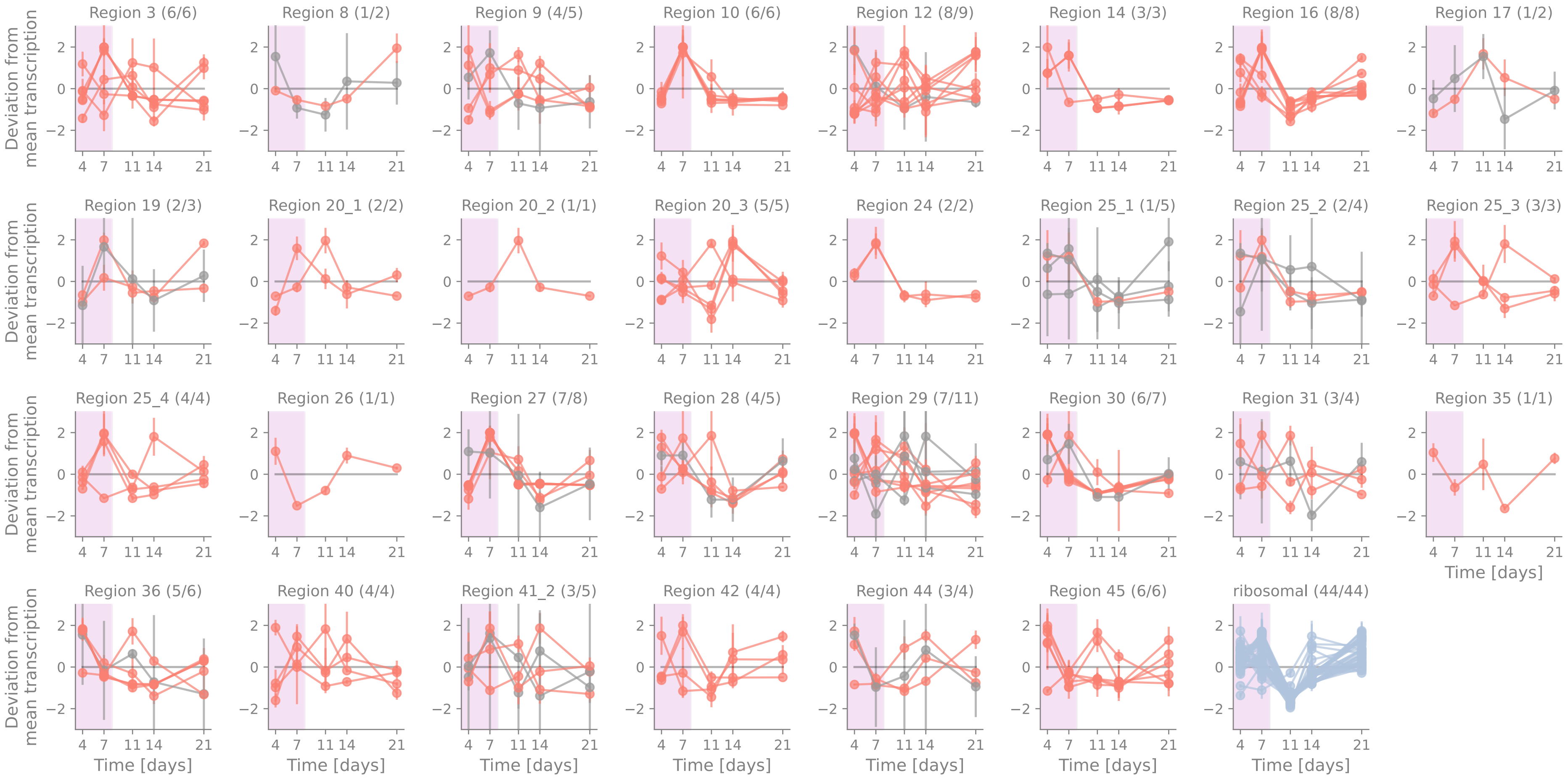


*Suppl. Fig. 6.* Temporal transcription of additional biosynthetic genes of NRP-, PK-related BGCs. Each subplot corresponds to one of the 25 antiSMASH-predicted BGC regions associated with the production of NRPs, PKs, or their hybrids. Some BGC regions contain several BGCs. We split these into the potential BGC subregions and represent each as an individual subplot indicated with underscores and subregion numbers. The numbers in brackets show the number of differentially transcribed additional biosynthetic genes and the total number of additional biosynthetic genes in the BGC (sub)region. The x-axis represents the growth time in days, the y-axis the deviation from mean transcription (z-score). Each curve corresponds to one additional biosynthetic gene. The data points show the mean over the replicates at the time point, the error bars the adjusted standard deviation, and the pink shading the exponential growth phase.

Differentially transcribed genes are represented in coral red, genes without a significant difference in transcription over the time-course experiment in grey. The last subplot shows the temporal transcription of 44 ribosomal genes as a comparison.

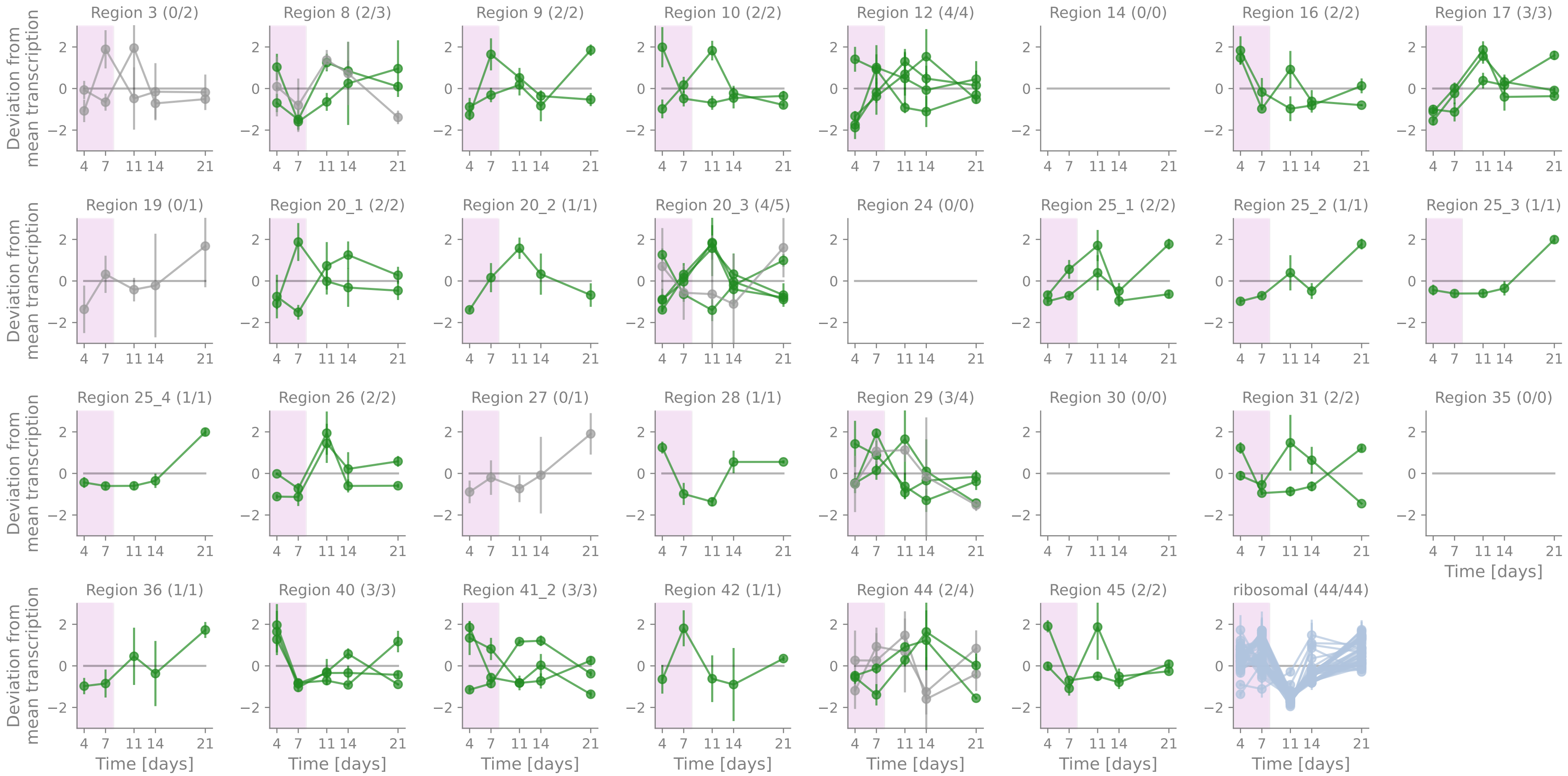


*Suppl. Fig. 7.* Temporal transcription of regulatory genes of NRP-, PK-related BGCs. Each subplot corresponds to one of the 25 antiSMASH-predicted BGC regions associated with the production of NRPs, PKs, or their hybrids. Some BGC regions contain several BGCs. We split these into the potential BGC subregions and represent each as an individual subplot indicated with underscores and subregion numbers. The numbers in brackets show the number of differentially transcribed regulatory genes and the total number of regulatory genes in the BGC (sub)region. The x-axis represents the growth time in days, the y-axis the deviation from mean transcription (z-score). Each curve corresponds to one regulatory gene. The data points show the mean over the replicates at the time point, the error bars the adjusted standard deviation, and the pink shading the exponential growth phase. Differentially transcribed genes are represented in green, genes without a significant difference in transcription over the time-course experiment in grey. The last subplot shows the temporal transcription of 44 ribosomal genes as a comparison.

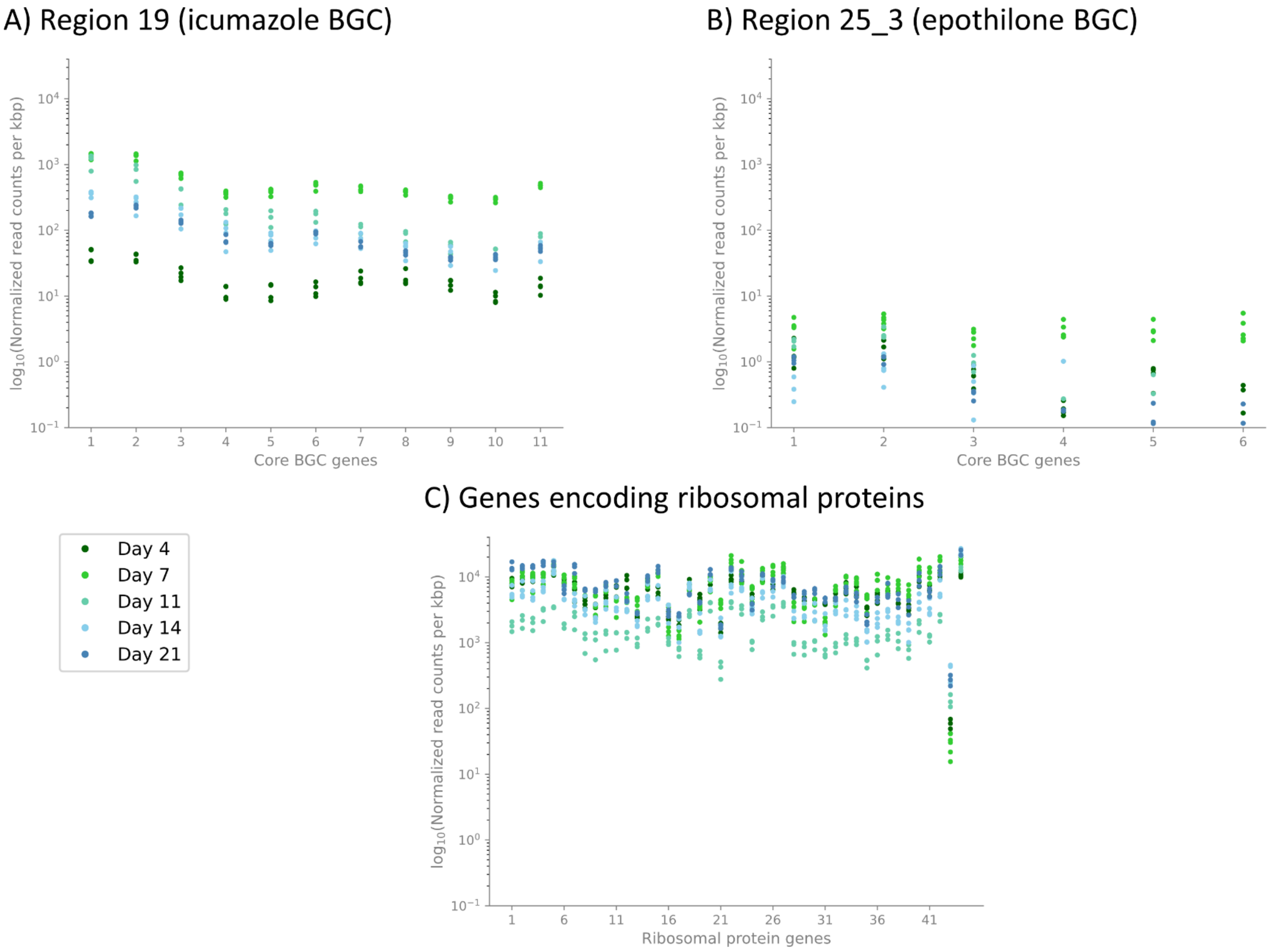
A) Region 19 (icumazole BGC)
B) Region 25_3 (epothilone BGC)
C) Genes encoding ribosomal proteins
log10(Normalized read counts per kbp)
Core BGC genes
Ribosomal protein genes
Day 4
Day 7
Day 11
Day 14
Day 21

*Suppl. Fig. 8.* Normalized read counts from DESeq2 of icumazole and epothilone BGC core genes, and ribosomal protein-related genes. The different colors represent the different growth days from day 4 in dark green, day 7 in light green, day 11 in turquoise, day 14 in light blue, and day 21 in dark blue. Each growth day was sampled with three to five replicates. **A)** Normalized read counts per kilo base pair of the BGC core genes associated with the production of icumazole. **B)** Normalized read counts per kilo base pair of the BGC core genes of BGC subregion 25_3 associated with the production of epothilone. **C)** Normalized read counts per kilo base pair of 44 genes encoding ribosomal proteins.

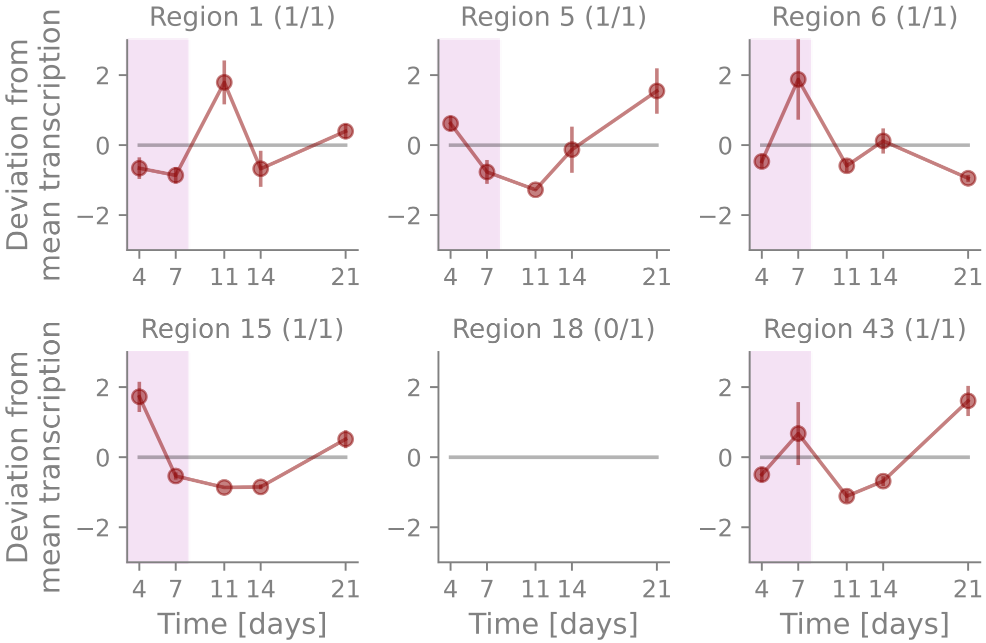


*Suppl. Fig. 9.* Temporal transcription of core genes of terpene-related BGCs. Each subplot corresponds to one of the six antiSMASH-predicted BGC regions associated with the production of terpenes. The numbers in brackets show the number of differentially transcribed core genes and the total number of core genes in the BGC (sub)region. The x-axis represents the growth time in days, the y-axis the deviation from mean transcription (z-score). Each curve corresponds to one core gene. The data points show the mean over the replicates at the time point, the error bars the adjusted standard deviation, and the pink shading the exponential growth phase. Differentially transcribed genes are represented in dark red, genes without a significant difference in transcription over the time-course experiment in grey. Genes that were filtered out by DESeq2 independent filtering are not shown (region 18).

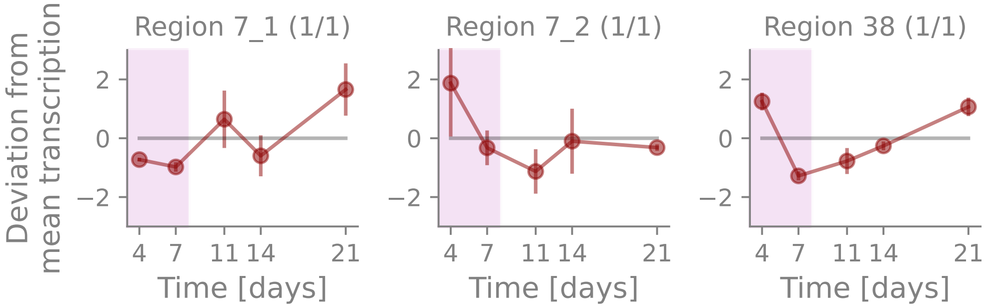


*Suppl. Fig. 10.* Temporal transcription of core genes of predicted BGCs associated with resorcinol, phenazine, or phosphonate production. The antiSMASH-predicted region 7 contains two potential BGCs. BGC subregion 7_1 might produce resorcinol, and subregion 7_2 phenazine. The antiSMASH-predicted BGC region 38 is associated with phosphonate production. The numbers in brackets show the number of differentially transcribed core genes and the total number of core genes in the BGC (sub)region. The x-axis represents the growth time in days, the y-axis the deviation from mean transcription (z-score). Each curve corresponds to one core gene. The data points show the mean over the replicates at the time point, the error bars the adjusted standard deviation, and the pink shading the exponential growth phase. Differentially transcribed genes are represented in dark red.

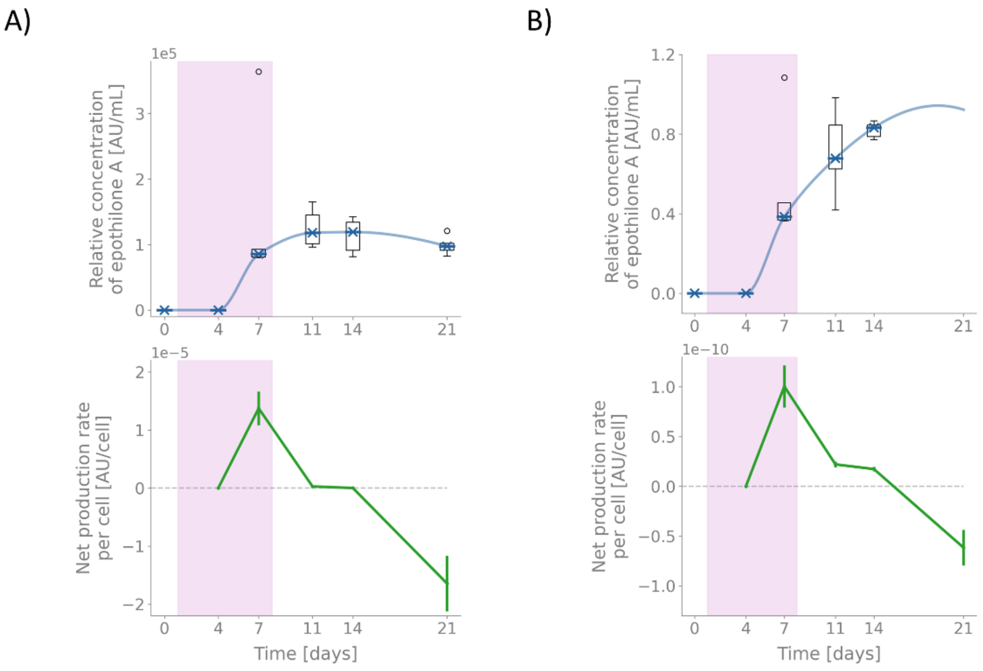


*Suppl. Fig. 11.* Production of epothilone A estimated from mass peak and UV absorption. **A)** is focused on the estimation of epothilone A based on the mass peak at 494.2571 and retention time around 9.05 min. **B)** shows the estimation of epothilone A based on the UV absorption at 250 nm and retention time of 9.05 min. The upper panels (blue) show the relative concentration of product. Represented are the box-plots over five replicates. The curves were generated by monotone piecewise cubic interpolation over the median values at each time point. The lower panels (green) show the production rate per cell based on the slope of the compound concentration per mL and the mean cell count per mL at a time point. The error bars correspond to the propagated error. The UV absorption on day 21 was ambiguous and thus excluded from the analysis. For the calculation of the net production rate per cell on day 21 in B, the estimated slope from the fitted curve of the relative concentration was used. The pink shading indicates the exponential growth phase.

*Suppl. Table S1.* BGC regions predicted by antiSMASH and additionally proposed subregions (the latter are indicated by the underscore in the region identifier and grey font). The + and - signs next to the number of genes indicates how many genes were located on each DNA strand.

| BGC region | Type | From | To | Most similar known BGC (MIBiG database) | Similarity[1] | core genes | additional genes | transport genes | regulatory genes | other genes | Median normalized read counts per kilobase (on day of maximal transcription) |
|---|---|---|---|---|---|---|---|---|---|---|---|
| 1 | terpene | 79,176 | 95,632 | | | 1 (-) | 3 (-) | 0 | 1 (+) | 3 (+), 3 (-) | 11 (day 11) |
| 2 | RiPP-like | 236,551 | 245,218 | | | 1 (+) | 0 | 0 | 0 | 3 (+), 6 (-) | 87 (day 11) |
| 3 | NRPS | 384,689 | 427,726 | | | 1 (-) | 2 (+), 4 (-) | 1 (-) | 2 (-) | 18 (+), 9 (-) | 263 (day 7) |
| 4 | RRE-containing, RiPP-like | 604,652 | 625,342 | | | 2 (+) | 2 (+), 2 (-) | 0 | 3 (+) | 4 (+), 4 (-) | 315 (day 7) |
| 5 | terpene | 889,356 | 909,328 | | | 1 (-) | 3 (+) | 0 | 2 (+), 1 (-) | 4 (+), 3 (-) | 315 (day 21) |
| 6 | terpene | 1,104,769 | 1,123,322 | carotenoid | 100% | 1 (+) | 3 (+), 1 (-) | 0 | 0 | 4 (+), 7 (-) | 273 (day 7) |
| 7 | resorcinol, phenazine | 1,319,935 | 1,367,558 | kirromycin | 5% | 2 (-) | 4 (+), 2 (-) | 1 (+), 1 (-) | 2 (+), 1 (-) | 10 (+), 11 (-) | |
| 7_1 | resorcinol | 1,319,935 | 1,357,109 | | | 1 (-) | 3 (+), 2 (-) | 1 (-) | 1 (+), 1 (-) | 8 (+), 10 (-) | 15 (day 21) |
| 7_2 | phenazine | 1,348,673 | 1,367,558 | | | 1 (-) | 1 (+) | 1 (+) | 1 (+) | 6 (+), 3 (-) | 14 (day 4) |
| 8 | hglE-KS | 1,622,851 | 1,676,650 | | | 2 (+) | 2 (+) | 3 (-) | 2 (+), 1 (-) | 6 (+), 15 (-) | 87 (day 21) |
| 9 | NRPS, T1PKS | 1,985,881 | 2,079,530 | nostocyclo peptide A2 | 28% | 3 (-) | 3 (+), 2 (-) | 2 (+) | 1 (+), 1 (-) | 11 (+), 7 (-) | 2 (day 4) |
| 10 | T1PKS | 2,462,829 | 2,559,365 | azalomycin F3a | 17% | 3 (+) | 1 (+), 5 (-) | 1 (+) | 2 (-) | 15(+), 5(-) | 126 (day 7) |

| | | | | | | | | | | | |
|---|---|---|---|---|---|---|---|---|---|---|---|
| 11 | RiPP-like | 2,642,544 | 2,652,117 | | | 1 (-) | 0 | 0 | 0 | 4 (+), 7 (-) | 32 (day 7) |
| 12 | NRPS | 2,745,866 | 2,792,667 | | | 1 (+) | 8 (+), 1 (-) | 0 | 2 (+), 2 (-) | 9 (+), 7 (-) | Not significantly differentially expressed |
| 13 | RiPP-like | 3,181,370 | 3,191,533 | | | 1 (+) | 1 (+) | 0 | 1 (-) | 6 (+) | Not significantly differentially expressed |
| 14 | T3PKS | 3,530,718 | 3,571,815 | alkylpyrone-407 / alkylpyrone-393 | 6% | 1 (-) | 1 (+), 2 (-) | 2 (+) | 0 | 18 (+), 7 (-) | 396 (day 7) |
| 15 | terpene | 3,715,607 | 3,736,715 | geosmin | 100% | 1 (+) | 0 | 2 (+) | 0 | 11 (+), 2 (-) | 35 (day 4) |
| 16 | NRPS | 3,998,980 | 4,047,658 | leinamycin | 4% | 1 (-) | 3 (+), 5 (-) | 1 (-) | 2 (-) | 4 (+), 8 (-) | 25 (day 7) |
| 17 | T1PKS | 4,742,391 | 4,800,650 | myxothiazol | 42% | 3 (+) | 1 (+), 1 (-) | 0 | 1 (+), 2 (-) | 6 (+), 7 (-) | 3 (day 11) |
| 18 | terpene | 5,252,143 | 5,271,158 | | | 1 (-) | 2 (+), 3 (-) | 0 | 0 | 6 (+), 2 (-) | Not significantly differentially expressed |
| 19 | T1PKS, NRPS | 5,491,866 | 5,600,251 | icumazole[2] | 100% | 11 (+) | 3 (+) | 0 | 1 (-) | 16 (+), 8 (-) | 427 (day 7) |
| 20 | NRPS, T1PKS | 5,648,140 | 5,752,549 | BE-43547A1 / BE-43547A2 / BE-43547B1 / BE-43547B2 / BE-43547B3 / BE-43547C1 / | 13% | 5 (+), 3 (-) | 5 (+), 2 (-) | 1 (-) | 3 (+), 4 (-) | 11 (+), 17 (-) | |

| | | | | | | | | | | | |
|---|---|---|---|---|---|---|---|---|---|---|---|
| | | | | BE-43547C2 | | | | | | | |
| 20_1 | NRPS | 5,648,140 | 5,674,889 | | | 1 (+) | 1 (+), 1 (-) | 0 | 2 (+) | 4 (+), 7 (-) | 5 (day 7) |
| 20_2 | NRPS, T1PKS | 5,671,779 | 5,703,491 | | | 3 (-) | 1 (+) | 1 (-) | 1 (-) | 1 (+), 3 (-) | 6 (day 4) |
| 20_3 | NRPS, T1PKS | 5,695,840 | 5,752,549 | | | 4 (+) | 4 (+), 1 (-) | 1 (-) | 1 (+), 4 (-) | 7 (+), 10 (-) | 16 (day 14) |
| 21 | thioamitides | 5,855,719 | 5,874,374 | | | 2 (+) | 2 (-) | 2 (-) | 0 | 6 (+), 7 (-) | 43 (day 11) |
| 22 | RiPP-like | 5,960,537 | 5,970,390 | | | 1 (-) | 1 (-) | 1 (-) | 0 | 4 (+), 4 (-) | 260 (day 4) |
| 23 | thioamitides | 6,123,337 | 6,144,000 | | | 2 (+) | 1 (+), 2 (-) | 0 | 1 (+), 1 (-) | 7 (+), 4 (-) | 33 (day 4) |
| 24 | NRPS, T1PKS | 6,595,997 | 6,657,138 | ajudazol A | 15% | 3 (+) | 2 (+) | 2 (+), 2 (-) | 0 | 12 (+), 15 (-) | 31 (day 7) |
| 25 | T1PKS, NRPS, transAT-PKS | 7,022,888 | 7,294,347 | epothilone | 100% | 18 (-) | 4 (+), 8 (-) | 4 (+), 3 (-) | 2 (+), 1 (-) | 32 (+), 59 (-) | |
| 25_1 | NRPS, T1PKS | 7,022,888 | 7,069,383 | | | 1 (-) | 2 (+), 3 (-) | 4 (+), 1 (-) | 1 (+), 1 (-) | 6 (+), 7 (-) | 491 (day 11) |
| 25_2 | transAT-PKS | 7,053,755 | 7,138,618 | | | 8 (-) | 1 (+), 3 (-) | 2 (+), 1 (-) | 1 (-) | 12 (+), 11 (-) | 16 (day 7) |
| 25_3 | NRPS, T1PKS | 7,138,217 | 7,231,683 | epothilone | 100% | 6 (-) | 1 (+), 2 (-) | 1 (-) | 1 (+) | 8 (+), 25 (-) | 3 (day 7) |
| 25_4 | NRPS | 7,215,071 | 7,294,347 | glidopeptin | 62% | 3 (-) | 1 (+), 3 (-) | 2 (-) | 1 (+) | 13 (+), 27 (-) | 72 (day 7) |
| 26 | T1PKS, NRPS | 7,370,296 | 7,417,399 | | | 2 (-) | 1 (+) | 0 | 2 (-) | 17 (+), 11 (-) | 8 (day 11) |
| 27 | T1PKS | 8,196,802 | 8,240,786 | LL-D49194α1 (LLD) | 5% | 1 (-) | 3 (+), 5 (-) | 0 | 1 (+) | 15 (+), 7 (-) | 504 (day 7) |

| | | | | | | | | | | | |
|---|---|---|---|---|---|---|---|---|---|---|---|
| 28 | NRPS | 8,287,155 | 8,333,743 | | | 1 (-) | 5 (-) | 1 (+) | 1 (+) | 12 (+), 8 (-) | 8 (day 7) |
| 29 | hglE-KS, T1PKS | 8,505,288 | 8,551,992 | | | 1 (-) | 3 (+), 8 (-) | 0 | 3 (+), 1 (-) | 11 (+), 6 (-) | 3 (day 4) |
| 30 | NRPS-like | 9,144,005 | 9,184,647 | syringomycin | 35% | 1 (+) | 6 (+), 1 (-) | 1 (+) | 0 | 16 (+), 8 (-) | 187 (day 14) |
| 31 | arylpolyene | 10,045,864 | 10,090,928 | N-tetradecanoyl tyrosine | 6% | 2 (-) | 3 (+), 1 (-) | 1 (-) | 1 (+), 1 (-) | 10 (+), 19 (-) | 59 (day 4) |
| 32 | LAP, RRE-containing | 10,176,352 | 10,200,167 | | | 3 (+) | 1 (-) | 0 | 3 (+) | 4 (+), 7 (-) | Not significantly differentially expressed |
| 33 | RiPP-like | 10,223,454 | 10,232,661 | | | 1 (+) | 0 | 0 | 1 (+) | 7 (+), 2 (-) | 55 (day 21) |
| 34 | thioamitides | 10,339,625 | 10,361,669 | | | 2 (+) | 0 | 0 | 2 (+) | 8 (+), 2 (-) | 9 (day 21) |
| 35 | T1PKS | 10,778,449 | 10,836,996 | jomthonic acid A / jomthonic acid B / jomthonic acid C | 5% | 2 (-) | 1 (+) | 1 (+) | 0 | 8 (+), 15 (-) | 196 (day 11) |
| 36 | NRPS | 11,005,691 | 11,051,523 | | | 1 (+) | 5 (+), 1 (-) | 1 (-) | 1 (+) | 11 (+), 9 (-) | 17 (day 4) |
| 37 | microviridin | 11,578,315 | 11,599,037 | | | 2 (+) | 1 (-) | 3 (-) | 0 | 4 (+), 3 (-) | 12674 (day 14) |
| 38 | phosphonate | 11,734,808 | 11,776,472 | | | 1 (+) | 2 (+), 2 (-) | 0 | 1 (+), 1 (-) | 14 (+), 11 (-) | 290 (day 4) |
| 39 | RiPP-like | 12,285,975 | 12,296,835 | | | 1 (+) | 0 | 0 | 0 | 4 (+), 4 (-) | 509 (day 21) |
| 40 | NRPS-like | 12,637,197 | 12,680,565 | | | 1 (+) | 2 (+), 2 (-) | 0 | 3 (+) | 13 (+), 18 (-) | 250 (day 4) |
| 41 | redox-cofactor, | 13,198,502 | 13,290,016 | leupyrrin | 17% | 8 (-) | 1 (+), 4 (-) | 3 (+), 2 (-) | 1 (+), 2 (-) | 14 (+), 17 (-) | |

| | | | | | | | | | | | |
|---|---|---|---|---|---|---|---|---|---|---|---|
| | NRPS, T1PKS | | | | | | | | | | |
| 41_1 | redox-cofactor | 13,198,502 | 13,220,204 | | | 2 (-) | 1 (-) | 2 (+), 1 (-) | 1 (-) | 5 (+), 8 (-) | 56 (day 4) |
| 41_2 | NRPS, T1PKS | 13,210,668 | 13,290,016 | | | 6 (-) | 1 (+), 4 (-) | 3 (+), 1 (-) | 1 (+), 2 (-) | 10 (+), 13 (-) | 2 (day 7) |
| 42 | NRPS-like, betalactone | 13,557,589 | 13,600,174 | rhizomide A / rhizomide B / rhizomide C | 100% | 2 (+) | 1 (+), 3 (-) | 7 (-) | 1 (-) | 9 (+), 8 (-) | 17 (day 7) |
| 43 | terpene | 13,674,072 | 13,695,043 | eremophilene | 100% | 1 (+) | 2 (+), 2 (-) | 1 (-) | 2 (-) | 2 (+), 1 (-) | 439 (day 21) |
| 44 | NRPS-like | 13,929,666 | 13,971,809 | | | 1 (-) | 4 (-) | 4 (-) | 3 (+), 1 (-) | 13 (+), 8 (-) | 74 (day 4) |
| 45 | NRPS, T1PKS | 14,457,907 | 14,535,411 | tubulysin A | 21% | 5 (+) | 2 (+), 4 (-) | 0 | 2 (-) | 9 (+), 11 (-) | 8 (day 4) |

[1] Please note that the similarity score indicates the percentage of genes in the MIBiG cluster that show amino acid similarity with genes in the predicted BGC. It does not consider the order of the genes in the cluster, nor does it indicate the percentage of amino acid similarity between matched genes.

[2] (Xie et al., 2022)

*Suppl. Table S2.* Over-/Under-representation analysis of NRP-, PK-related BGC core genes among transcription clusters

| Transcription cluster | Genes in cluster | BGC core genes in cluster | over- / under-represented | adjusted p-value |
|---|---|---|---|---|
| Cluster 1 | 985 | 3 | under-represented | 4.36e-2 |
| Cluster 2 | 2,857 | 18 | / | / |
| Cluster 3 | 395 | 2 | / | / |
| Cluster 4 | 1,423 | 40 | over-represented | <6.27e-11 |
| Cluster 5 | 2,669 | 12 | under-represented | 6.34e-3 |

*Suppl. Table S3.* Genes encoding ribosomal proteins

| Locus name | Gene name | Product |
|---|---|---|
| NQZ70_00662 | rplK | 50S ribosomal protein L11 |
| NQZ70_00663 | rplA | 50S ribosomal protein L1 |
| NQZ70_00664 | rplJ | 50S ribosomal protein L10 |
| NQZ70_00665 | rplL | 50S ribosomal protein L7/L12 |
| NQZ70_01256 | rpsL | 30S ribosomal protein S12 |
| NQZ70_01257 | rpsG | 30S ribosomal protein S7 |
| NQZ70_01260 | rpsJ | 30S ribosomal protein S10 |
| NQZ70_01261 | rplD | 50S ribosomal protein L4 |
| NQZ70_01262 | rplW | 50S ribosomal protein L23 |
| NQZ70_01263 | rplB | 50S ribosomal protein L2 |
| NQZ70_01264 | rpsS | 30S ribosomal protein S19 |
| NQZ70_02244 | | ribosomal protein S21 |
| NQZ70_02460 | rplC | 50S ribosomal protein L3 |
| NQZ70_03029 | rpmGA | 50S ribosomal protein L33 1 |
| NQZ70_03140 | rplS | 50S ribosomal protein L19 |
| NQZ70_03144 | rpmH | 50S ribosomal protein L34 |
| NQZ70_04777 | rpmF | 50S ribosomal protein L32 |
| NQZ70_05741 | rpsP | 30S ribosomal protein S16 |
| NQZ70_05855 | rpsO | 30S ribosomal protein S15 |
| NQZ70_08229 | rpsA | 30S ribosomal protein S1 |
| NQZ70_08293 | rplI | 50S ribosomal protein L9 |
| NQZ70_08597 | rplT | 50S ribosomal protein L20 |
| NQZ70_08598 | rpmI | 50S ribosomal protein L35 |
| NQZ70_08772 | rpsI | 30S ribosomal protein S9 |
| NQZ70_08773 | rplM | 50S ribosomal protein L13 |
| NQZ70_09247 | rpmA | 50S ribosomal protein L27 |
| NQZ70_09248 | rplU | 50S ribosomal protein L21 |
| NQZ70_09263 | rpsD | 30S ribosomal protein S4 |
| NQZ70_09264 | rpsK | 30S ribosomal protein S11 |
| NQZ70_09265 | rpsM | 30S ribosomal protein S13 |
| NQZ70_09266 | rpmJ | 50S ribosomal protein L36 |
| NQZ70_09270 | rplO | 50S ribosomal protein L15 |
| NQZ70_09273 | rplR | 50S ribosomal protein L18 |
| NQZ70_09274 | rplF | 50S ribosomal protein L6 |
| NQZ70_09275 | rpsH | 30S ribosomal protein S8 |
| NQZ70_09276 | rpsZ | 30S ribosomal protein S14 type Z |
| NQZ70_09278 | rplX | 50S ribosomal protein L24 |
| NQZ70_09279 | rplN | 50S ribosomal protein L14 |
| NQZ70_09281 | rpmC | 50S ribosomal protein L29 |
| NQZ70_09282 | rplP | 50S ribosomal protein L16 |
| NQZ70_09283 | rpsC | 30S ribosomal protein S3 |
| NQZ70_09284 | rplV | 50S ribosomal protein L22 |
| NQZ70_09519 | rpmG | 50S ribosomal protein L33 |
| NQZ70_10413 | rpmE | 50S ribosomal protein L31 |

*Suppl. Table S4.* COG categories and their description. Categories in grey are not applied to prokaryotes.

| **COG category** | **Description** |
|---|---|
| J | Translation, ribosomal structure and biogenesis |
| A | RNA processing and modification |
| K | Transcription |
| L | Replication, recombination and repair |
| B | Chromatin structure and dynamics |
| D | Cell cycle control, cell division, chromosome partitioning |
| Y | Nuclear structure |
| V | Defense mechanisms |
| T | Signal transduction mechanisms |
| M | Cell wall/membrane/envelope biogenesis |
| N | Cell motility |
| Z | Cytoskeleton |
| W | Extracellular structures |
| U | Intracellular trafficking, secretion, and vesicular transport |
| O | Posttranslational modification, protein turnover, chaperones |
| X | Mobilome: prophages, transposons |
| C | Energy production and conversion |
| G | Carbohydrate transport and metabolism |
| E | Amino acid transport and metabolism |
| F | Nucleotide transport and metabolism |
| H | Coenzyme transport and metabolism |
| I | Lipid transport and metabolism |
| P | Inorganic ion transport and metabolism |
| Q | Secondary metabolites biosynthesis, transport and catabolism |
| R | General function prediction only |
| S | Function unknown |

*Suppl. Table S5.* Functional analysis of transcription clusters.

| **Transcription cluster** | **annotated/all genes in cluster** | **over-/under-represented** | **COG category** | **Description** | **differentially expressed genes in COG** | **COG genes in cluster** | **adjusted p-value** |
|---|---|---|---|---|---|---|---|
| Cluster 1 | 713/985 | over-represented | G | Carbohydrate transport and metabolism | 409 | 72 | 3.16e-3 |
| Cluster 2 | 2,132/2,857 | over-represented | M | Cell wall/membrane/envelope biogenesis | 401 | 170 | 8.35e-3 |
| | | over-represented | F | Nucleotide transport and metabolism | 133 | 67 | 2.98e-3 |
| Cluster 3 | 272/395 | / | / | / | / | / | / |
| Cluster 4 | 1,104/1,423 | over-represented | P | Inorganic ion transport and metabolism | 290 | 76 | 3.49e-3 |
| | | over-represented | J | Translation, ribosomal structure and biogenesis | 222 | 57 | 1.60e-2 |
| | | over-represented | Q | Secondary metabolites biosynthesis, transport and catabolism | 339 | 86 | 3.49e-3 |

| | | | | | | | |
|---|---|---|---|---|---|---|---|
| Cluster 5 | 1,860/2,669 | under-represented | E | Amino acid transport and metabolism | 364 | 88 | 1.26e-2 |
| | | under-represented | P | Inorganic ion transport and metabolism | 290 | 67 | 1.26e-2 |
| | | under-represented | J | Translation, ribosomal structure and biogenesis | 222 | 40 | 1.82e-4 |
| | | under-represented | M | Cell wall/membrane/envelope biogenesis | 401 | 101 | 2.29e-2 |
| | | under-represented | H | Coenzyme transport and metabolism | 249 | 57 | 1.26e-2 |
| | | under-represented | F | Nucleotide transport and metabolism | 133 | 25 | 1.14e-2 |
| | | over-represented | O | Posttranslational modification, protein turnover, chaperones | 271 | 108 | 9.71e-3 |
